# PITPβ promotes COPI vesicle fission through lipid transfer and membrane contact formation

**DOI:** 10.1101/2024.05.27.596058

**Authors:** Kunyou Park, Sungeun Ju, Hyewon Choi, Peng Gao, Geul Bang, Jung Hoon Choi, Jiwon Jang, Andrew J. Morris, Byung-Ho Kang, Victor W. Hsu, Seung-Yeol Park

**Affiliations:** Department of Life Sciences, Pohang University of Science and Technology, Pohang, Gyeongbuk 37673, Republic of Korea; Division of Rheumatology, Inflammation and Immunity, Brigham and Women’s Hospital, and Department of Medicine, Harvard Medical School, Boston, MA 02115, USA; School of Life Sciences, State Key Laboratory of Agrobiotechnology, The Chinese University of Hong Kong, Shatin, New Territories, Hong Kong, China; Research Center for Bioconvergence Analysis, Korea Basic Science Institute, Cheongju, Republic of Korea; Department of Bio-Chemical Analysis, Korea Basic Science Institute, Cheongju, Republic of Korea; University of Arkansas for Medical Sciences and Central Arkansas Veterans Affairs Healthcare System, Little Rock, AR 72205, USA

**Keywords:** COPI vesicle formation, Golgi transport, Membrane contact, Lipid transfer

## Abstract

Intracellular transport among organellar compartments occurs in two general ways, by membrane-bound carriers or membrane contacts. Specific circumstances that involve the coordination of these two modes of transport remain to be defined. Studying Coat Protein I (COPI) transport, we find that phosphatidylcholine with short acyl chains (sPC) is delivered through membrane contact from the endoplasmic reticulum (ER) to sites of COPI vesicle formation at the Golgi to support the fission stage. Phosphatidylinositol transfer protein beta (PITPβ) plays a key role in this process, with the elucidation of this role advancing a new understanding of how PITPβ acts, providing a mechanistic understanding of a specific circumstance when vesicular transport requires membrane contact, and contributing to a basic understanding of how transport carriers in a model intracellular pathway are formed.

**Summary:** Specific circumstances that membrane contact is needed for vesicular transport remain to be defined. We find that a critical lipid is delivered through membrane contact to support the fission stage of a model intracellular transport pathway.

## Introduction

Transport by membrane-bound carriers occurs through conserved steps that are mediated by major classes of effectors. Coat complexes initiate transport by bending membrane to generate carriers and binding to cargoes for their sorting into these carriers. Subsequently, tether complexes act by docking the transport carriers to their target compartments. Soluble NEM-sensitive associated protein (SNAP) receptors (SNAREs) then complete transport by promoting the fusion of transport carriers with their target compartments. A detailed understanding of how these core transport effectors act, and how they are regulated, is being achieved for model pathways (Brocker et al., 2010; Cai et al., 2007; Donaldson and Jackson, 2011; McMahon and Boucrot, 2011; Miller and Barlowe, 2010; Mizuno-Yamasaki et al., 2012; Pucadyil and Schmid, 2009; Wickner and Schekman, 2008).

How transport occurs through membrane contacts is also being elucidated. This mode of transport has been found to play widespread roles in the delivery of lipids among organellar compartments. Multiple classes of proteins have been identified to possess lipid transfer activity. These lipid transfer proteins can also participate in contact formation. How these two functions are integrated to achieve lipid transport through membrane contact is also being elucidated (Antonny et al., 2018; Cohen et al., 2018; Lujan et al., 2021; Prinz et al., 2020; Reinisch and Prinz, 2021; Saheki and De Camilli, 2017; Venditti et al., 2020; Wong et al., 2019; Wu et al., 2018).

The ER forms contact with multiple organellar compartments through vesicle-associated membrane protein (VAMP)-associated proteins (VAPs) that reside on the ER membrane (Murphy and Levine, 2016). In the case of cholesterol transfer, VAP-A has been found to interact with oxysterol binding protein (OSBP) on the Golgi membrane to form contact between the ER and the trans-Golgi network (TGN). OSBP also possesses sterol transfer activity, and this activity is promoted by the counter-transfer of phosphatidylinositol 4-phosphate (PI4P) by OSBP. Through reconstitution studies, a mechanistic understanding of how OSBP couples its functions in lipid transfer and contact formation is being achieved (Mesmin et al., 2013).

Studies on vesicle formation by the Coat Protein I (COPI) complex have been contributing to a fundamental understanding of how transport carriers are generated (Hsu et al., 2009; Jackson, 2014; Lippincott-Schwartz and Liu, 2006; Pucadyil and Schmid, 2009). COPI vesicles act in retrograde transport at the Golgi complex and also from the Golgi to the endoplasmic reticulum (ER) (Brandizzi and Barlowe, 2013; Lee et al., 2004; Suda et al., 2017). Early studies identified coatomer, a multimeric complex, as the core components of the COPI complex, and ADP-Ribosylation Factor 1 (ARF1) as the small GTPase that regulates the recruitment of coatomer from the cytosol to the Golgi membrane to initiate COPI vesicle formation (Donaldson et al., 1992; Malhotra et al., 1989; Serafini et al., 1991; Waters et al., 1992). Subsequently, a GTPase activating protein (GAP) that deactivates ARF1 (known as ARFGAP1) was identified to act not only as its regulator (Cukierman et al., 1995), but also as an effector (Yang et al., 2002), in COPI vesicle formation.

The late stage of vesicle formation involves membrane fission, which severs the neck of coated buds to release them as coated vesicles. Brefeldin-A ADP-Ribosylation Substrate (BARS) has been identified to promote COPI vesicle fission (Yang et al., 2005). More recently, lipid-based mechanisms have also been discovered to play a key role (Park et al., 2019; Yang et al., 2008; Yang et al., 2011). COPI vesicle fission has been found to require the sequential actions of phospholipase D2 (PLD2) activity, which converts phosphatidylcholine (PC) to phosphatidic acid (PA), followed by lipid phosphate phosphatase 3 (LPP3) activity, which converts PA to diacylglycerol (DAG) (Park et al., 2019). The critical roles of PA and DAG require these lipids having short acyl chains, which facilitate the generation of extreme negative membrane curvature needed to drive vesicle fission to completion (Park et al., 2019).

A key question also arises from this recent elucidation. As short PC (sPC) is the precursor of short PA and DAG that are needed for COPI vesicle fission (Park et al., 2019), and PC synthesis occurs in the ER (Vance and Vance, 2004), how is sPC delivered from the ER to sites of COPI vesicle formation at the Golgi to support the fission stage? In this study, we find that PITPβ plays a key role in delivering sPC from the ER to sites of COPI vesicle formation at the Golgi to promote the fission stage. By combining cell-based and reconstitution approaches, we achieve a high level of spatial and temporal resolution in explaining how this delivery occurs. Our results define a specific circumstance that membrane contact is needed for vesicular transport, and advance a new understanding of how PITPβ acts. Moreover, as COPI transport has been a model for mechanistic studies on vesicle formation, our findings advance a basic understanding of how intracellular transport carriers are formed.

## Results

### COPI transport requires membrane contact between the ER and the Golgi

As lipid can be transferred among intracellular membrane compartments by transport carriers or membrane contacts, we initially sought to determine which mechanism is required for sPC delivery to support COPI vesicle fission. The COPII complex generates transport carriers from the ER for transport to the Golgi (Brandizzi and Barlowe, 2013; Lee et al., 2004; Suda et al., 2017). Prolonged inhibition of COPII transport disrupts the Golgi complex (Ward et al., 2001). Thus, we sought to inhibit COPII transport for short duration by treating cells pharmacologically with H89, which inhibits COPII transport while allowing the Golgi to remain intact (Lee and Linstedt, 2000). We first confirmed that this treatment inhibited COPII transport, as tracked by the transport of a model secretory protein, VSVG, from the ER to the Golgi (Fig. S1A). We also confirmed that the dose of H89 treatment allowed the Golgi to remain intact (Fig. S1B). We then found that COPI transport was not inhibited by this treatment, as tracked by the transport of a model COPI cargo, VSVG-KDELR, as previously described (Park et al., 2019; Yang et al., 2008; Yang et al., 2018; Yang et al., 2005) (Fig. 1A).

**Fig. 1.**
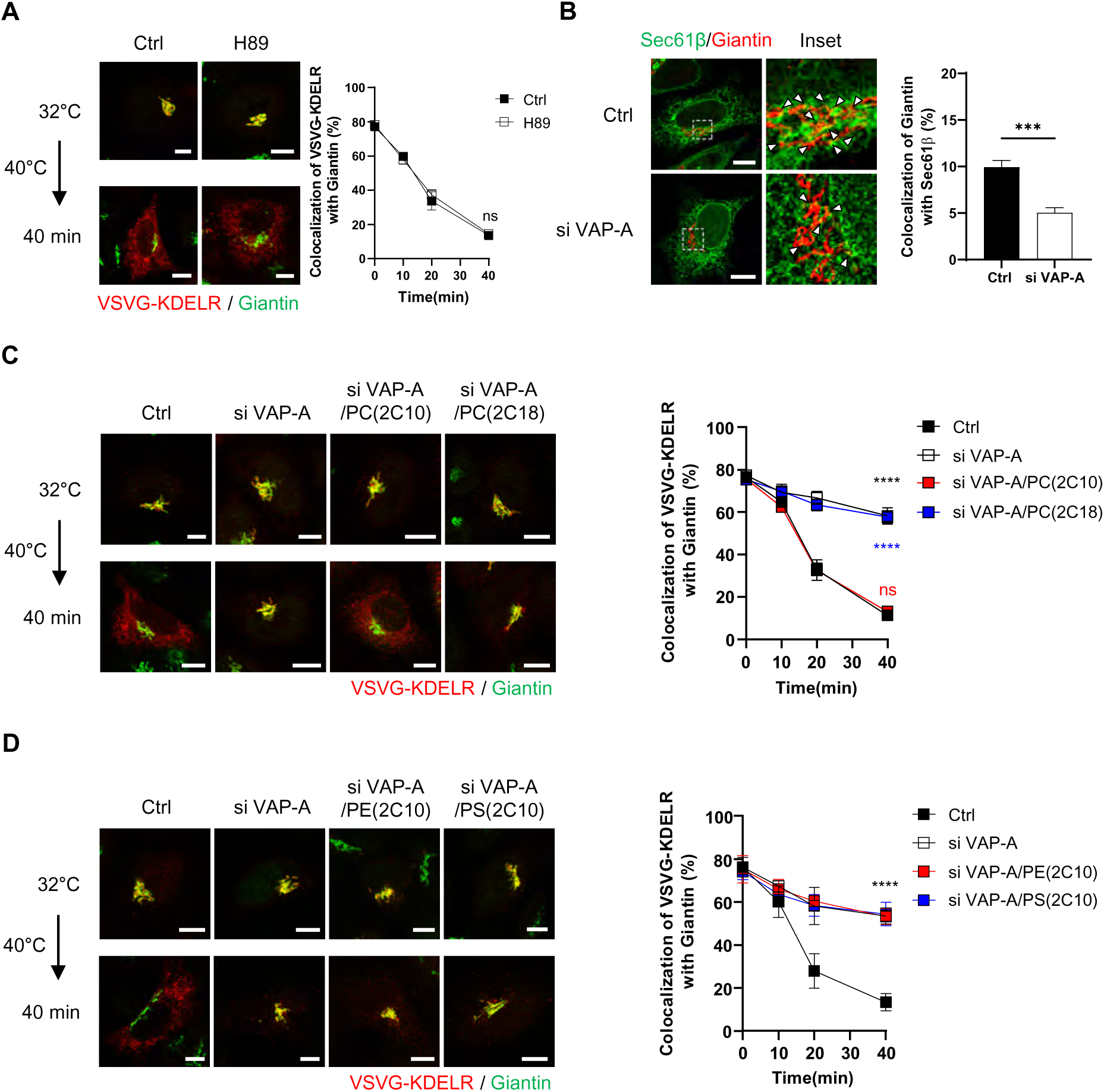
COPI transport requires membrane contact. Quantitative data are shown as mean ± s.e.m., with the number of independent experiments indicated. Statistics was performed using the two-tailed student t-test: **** p<0.0001, *** p<0.001, ns (non-significant) p>0.05. **(A)** COPI transport assay assessing the effect of 20 µM H89 treatment. Quantitation is shown on right, n=3. Representative confocal images are shown on left, VSVG-KDELR (red) and giantin (green), bar=10 µm. **(B)** Airyscan confocal microscopy examining the effect of siRNA against VAP-A on the colocalization of an ER marker (Sec61β) and a Golgi marker (giantin). Quantitation is shown on right, n=3. Representative images are shown on left, Sec61β (green), giantin (red), bar=10 µm. **(C)** COPI transport assay assessing the effect of siRNA against VAP-A followed by functional lipid rescue by feeding cells with different lengths of PC. Representative confocal images are shown on left, VSVG-KDELR (red), giantin (green), bar=10 µm. Quantitation is shown on right, n=4. **(D)** COPI transport assay assessing the effect of siRNA against VAP-A followed by functional lipid rescue by feeding cells with short forms of PE or PS. Quantitation is shown on right, n=3. Representative confocal images are shown on left, VSVG-KDELR (red) and giantin (green), bar=10 µm.

In light of the above result, we next examined whether membrane contact between the ER and the Golgi would be required COPI transport. The ER forms contacts with other organelles through VAMP-associated proteins (VAPs) expressed on the ER membrane (Murphy and Levine, 2016). As VAP-A has been found previously to participate in forming contact between the ER and the Golgi (Mesmin et al., 2013), we initially confirmed by confocal microscopy that a population of VAP-A colocalized with both an ER marker (Sec61β) and a Golgi marker (giantin) (Fig. S1C). We also confirmed that siRNA against VAP-A (Fig. S1D), using a sequence whose specificity had been previously documented (Nthiga et al., 2020), reduced the colocalization of these two markers (Fig. 1B). We then found that siRNA against VAP-A inhibited COPI transport (Fig. S1E).

To gain insight into whether this inhibition could be attributed to impaired sPC delivery, we next pursued a functional rescue approach. Lipids can be delivered into cells by incubating them in the culture medium in the presence of albumin (Pinot et al., 2014). We had previously taken this approach to deliver specific lipids to the Golgi (Park et al., 2019). We found that the inhibition of COPI transport induced by siRNA against VAP-A was rescued by feeding cells with PC having shorter (2C10), but not longer (2C18), acyl chains (Fig. 1C). We also confirmed lipid delivery to the Golgi by tracking a fluorescence-labeled PC fed to cells (Fig. S1F). Furthermore, we found that feeding cells with other major lipids of the Golgi in their shorter form (2C10), such as phosphatidylethanolamine (PE) or phosphatidylserine (PS), did not revert the inhibition in COPI transport (Fig. 1D). Thus, the collective results suggested that sPC delivery needed for COPI transport occurs through membrane contact rather than through COPII transport.

### PC transfer by PITPβ promotes COPI transport

As further confirmation, we next sought to identify a lipid transfer protein predicted to be involved. Phosphatidylinositol transfer protein beta (PITPβ) possesses PC transfer activity (Allen-Baume et al., 2002), and has been found previously to act in COPI transport (Carvou et al., 2010). However, a mechanistic understanding of how it promotes COPI transport had not been achieved. We first confirmed that siRNA against PITPβ (Fig. 2A) inhibited COPI transport (Fig. 2B). We also documented the targeting specificity of this inhibition with a rescue experiment (Fig. 2B). Moreover, the catalytic activity of PITPβ was needed, as a catalytic-dead form (C94A) could not rescue the inhibition (Fig. 2B). As controls, we found that this siRNA treatment did not affect anterograde transport into (Fig. 2C) and out of the Golgi (Fig. 2D), as tracked through the itinerary of VSVG. Thus, PITPβ showed specificity in supporting COPI transport.

**Fig. 2.**
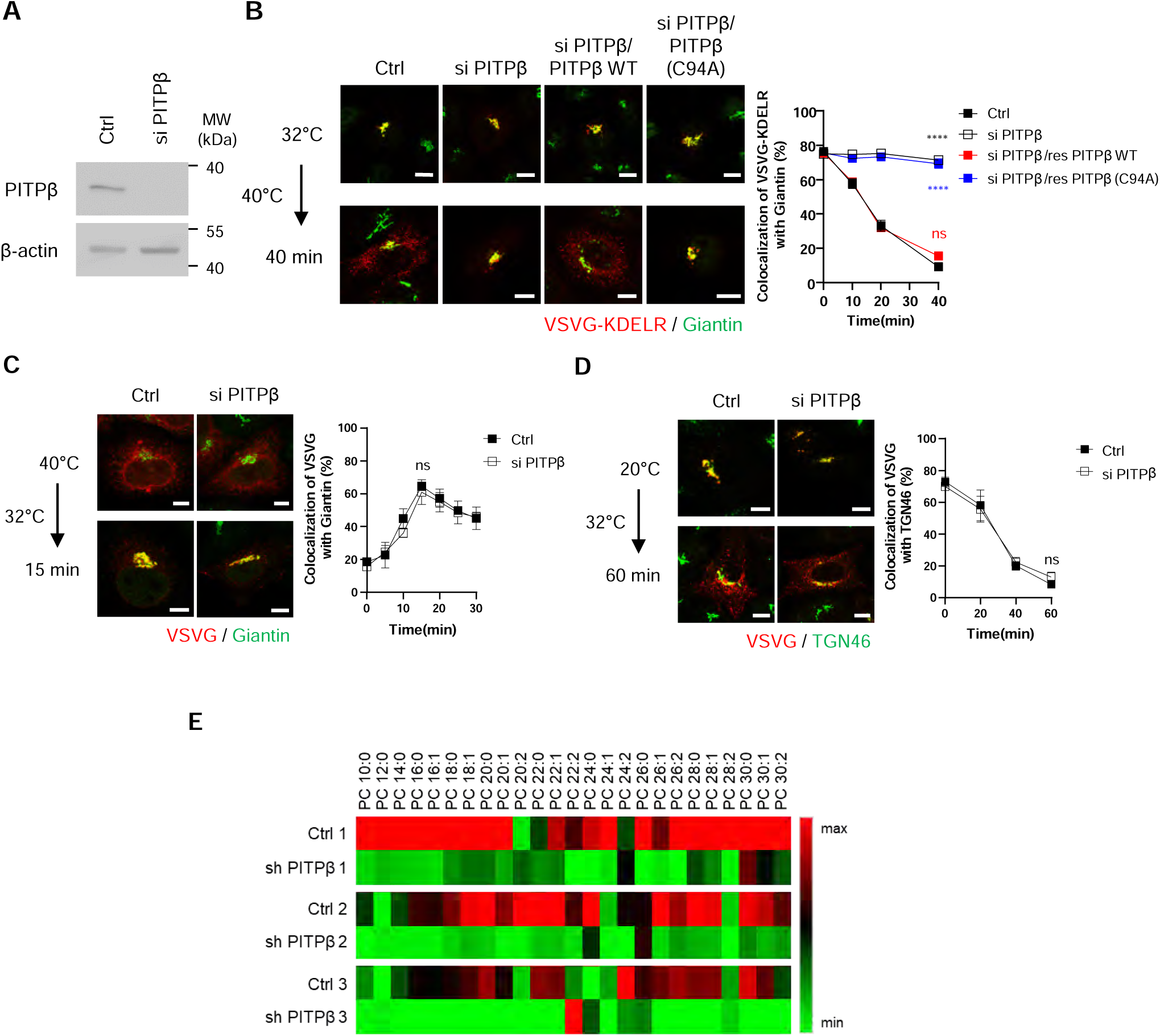
Characterizing the effects of siRNA against PITPβ. Quantitative data are shown as mean ± s.e.m, with the number of independent experiments indicated. Statistics was performed using the two-tailed student t-test: **** p<0.0001, ns p>0.05. **(A)** Efficacy of siRNA against PITPβ assessed by western blotting of whole cell lysate. β-actin level confirmed similar levels of lysate examined. **(B)** COPI transport assay measuring the effect of siRNA against PITPβ and rescue using either wild-type or catalytic dead (C94A) form. Representative confocal images are shown on left, VSVG-KDELR (red) and giantin (green), bar=10 µm. Quantitation is shown on right, n=7. **(C)** Transport assay assessing the effect of PITPβ depletion on the arrival of a model secretory cargo, VSVG, to the Golgi from the ER. Quantitation is shown on right, n=3. Representative confocal images are shown on left, VSVG (red) and giantin (green), bar=10 µm. **(D)** Transport assay assessing the effect of PITPβ depletion on the transport of VSVG from the Golgi to the plasma membrane. Quantitation is shown on right, n=3. Representative confocal images are shown on left, VSVG (red) and TGN46 (green), bar=10 µm. **(E)** Reduction of sPC levels in Golgi membrane upon PITPβ depletion as assessed by targeted lipidomics, n=3. Results are shown as heat map. PC species are categorized based on their total acyl chain length and unsaturations.

We next collected Golgi membrane from PITPβ-depleted cells to analyze PC level. Performing lipidomics, we confirmed that the levels of many PC species were reduced (Fig. S2). However, the conventional approach of lipidomics that we had undertaken, which is untargeted, had difficulty in detecting sPC species, having 28 total carbons or less. As these sPC species are less abundant than the longer PCs, we next developed a targeted approach, which took into consideration that shorter PCs are less hydrophobic than longer PCs due to shorter acyl chains. Taking this modified approach, we were able to detect sPC species and confirm that they were reduced in Golgi membrane isolated from cells treated with siRNA against PITPβ (Fig. 2E).

### PITPβ also acts in contact formation between the ER and COPI buds at the Golgi

Besides their catalytic activity, lipid transfer proteins have been found also to participate in forming membrane contacts (Antonny et al., 2018; Cockcroft and Raghu, 2018; Cohen et al., 2018; Prinz et al., 2020; Reinisch and Prinz, 2021; Venditti et al., 2020; Wong et al., 2019; Wu et al., 2018). Thus, to determine whether PITPβ possesses this additional capability, we initially performed confocal microscopy using Airyscan, which provides two-fold greater resolution than the conventional confocal approach. We detected a population of PITPβ colocalizing with both Sec61β (ER marker) and giantin (Golgi marker) (Fig. 3A). Quantitation revealed a low level of colocalization (Fig. 3B), which was expected, as colocalization should only occur for regions of the ER and the Golgi that form membrane contact. For this fraction, we found that siRNA against PITPβ reduced its level (Fig. 3B). As control, we found that siRNA against PITPβ did not reduce membrane contact between the ER and the trans-Golgi network (TGN), as tracked through the colocalization of Sec61β (marking the ER) and TGN46 (marking the TGN) (Fig 3C).

**Fig. 3.**
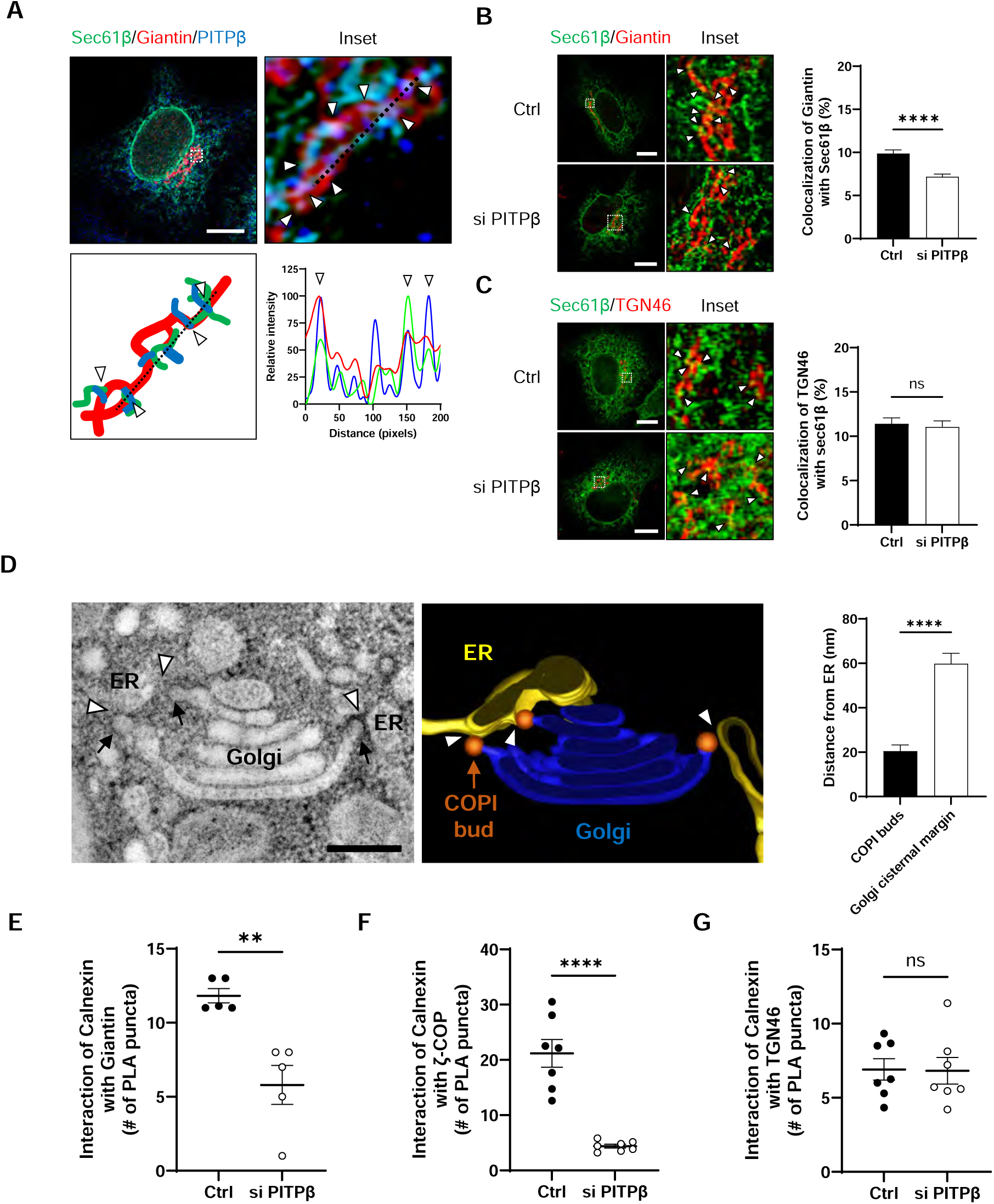
PITPβ promotes contact between the ER and COPI buds on the Golgi. Quantitative data are shown as mean ± s.e.m, with the number of independent experiments indicated. Statistics was performed using the two-tailed student t-test: **** p<0.0001, ** p<0.01, ns p>0.05. **(A)** Colocalization of PITPβ with ER marker (Sec61β) and Golgi marker (giantin) as assessed by confocal microscopy using Airyscan, PITPβ (blue), Sec61β (green), giantin (red). A representative image is shown in upper left panel, bar=10 μm. An inset is shown in upper right panel with arrow heads highlighting PITPβ colocalizing with both Sec61β and Giantin. A reconstruction of the region highlighted by the arrow heads in the inset is shown in lower left panel. Line scan along the dotted line in the inset is shown in the lower right panel. **(B)** The effect of siRNA against PITPβ on the colocalization of an ER marker (Sec61β) and a Golgi marker (giantin) as assessed by confocal microscopy using Airyscan. Quantitation is shown on right, n=6. Representative images are shown on left, bar=10 µm. Arrow heads highlight areas of colocalization. **(C)** Airyscan confocal microscopy examining the effect of siRNA against PITPβ on the colocalization of an ER marker (Sec61β) and a TGN marker (TGN46). Representative images are shown on left, bar=10 µm. Arrow heads highlight areas of colocalization. Quantitation is shown on right, n=4. **(D)** EM tomography showing COPI buds at the Golgi in close proximity to ER membrane. A representative tomographic image slice is shown in left panel with arrow heads pointing to COPI buds, bar=200 nm. 3D reconstruction of the Golgi, COPI buds, and ER elements is shown in right panel. Quantitation is shown on extreme right, comparing distance between COPI buds and ER membranes versus distance between Golgi cisternal margins and ER membranes, n=3. **(E)** Proximity ligation assay examining the effect of siRNA against PITPβ on the proximity between calnexin and giantin, n=5. **(F)** Proximity ligation assay assessing the effect of PITPβ depletion on the colocalization of calnexin and ζ-COP, n=5. **(G)** Proximity ligation assay examining the effect of siRNA against PITPβ on the proximity between calnexin and TGN46, n=4.

We next sought to detect the observed membrane contact at even higher resolution by pursuing electron tomography. We found that COPI buds were in closer proximity to ER membranes than the rest of the Golgi membrane (Fig. 3D). As this finding suggested that the ER could be contacting the Golgi more specifically at sites of COPI vesicle formation, we next sought further support by pursuing proximity ligation analysis. This approach provides greater resolution (detecting proteins within 40 nm of each other) than standard confocal microscopy (detecting proteins up to 200 nm from each other). We first found that siRNA against PITPβ reduced the proximity between calnexin (another ER marker) and giantin (Fig. 3E). We then tracked COPI buds through ζ-COP and found that siRNA against PITPβ reduced the proximity between calnexin and ζ-COP even more dramatically (Fig. 3F). As control, siRNA against PITPβ did not reduce the proximity between calnexin and TGN46 (Fig. 3G). Thus, these findings suggested that PITPβ acts in contact formation between the ER and the Golgi, with the latter more specifically at sites of COPI buds.

To further characterize how PITPβ acts in this role, we next performed a co-precipitation experiment and found that PITPβ associated with VAP-A in cells, and as control, PITPβ had no appreciable interaction with VAP-B (Fig. 4A). We then considered that VAPs have been found to interact with binding partners by recognizing an FFAT-like sequence on these partners (Murphy and Levine, 2016). As a sequence in PITPβ conforms to this motif, we mutated this sequence (F107A/F108A) and found that VAP-A could no longer interact directly with PITPβ (Fig. 4B). We also found that replacing the endogenous PITPβ with the mutant PITPβ inhibited COPI transport in cells (Fig. 4C). This replacement was achieved by performing siRNA against PITPβ followed by rescue using the mutant form. Thus, the results further confirmed a new function for PITPβ, acting to form membrane contact by interacting with VAP-A on the ER membrane.

**Fig. 4.**
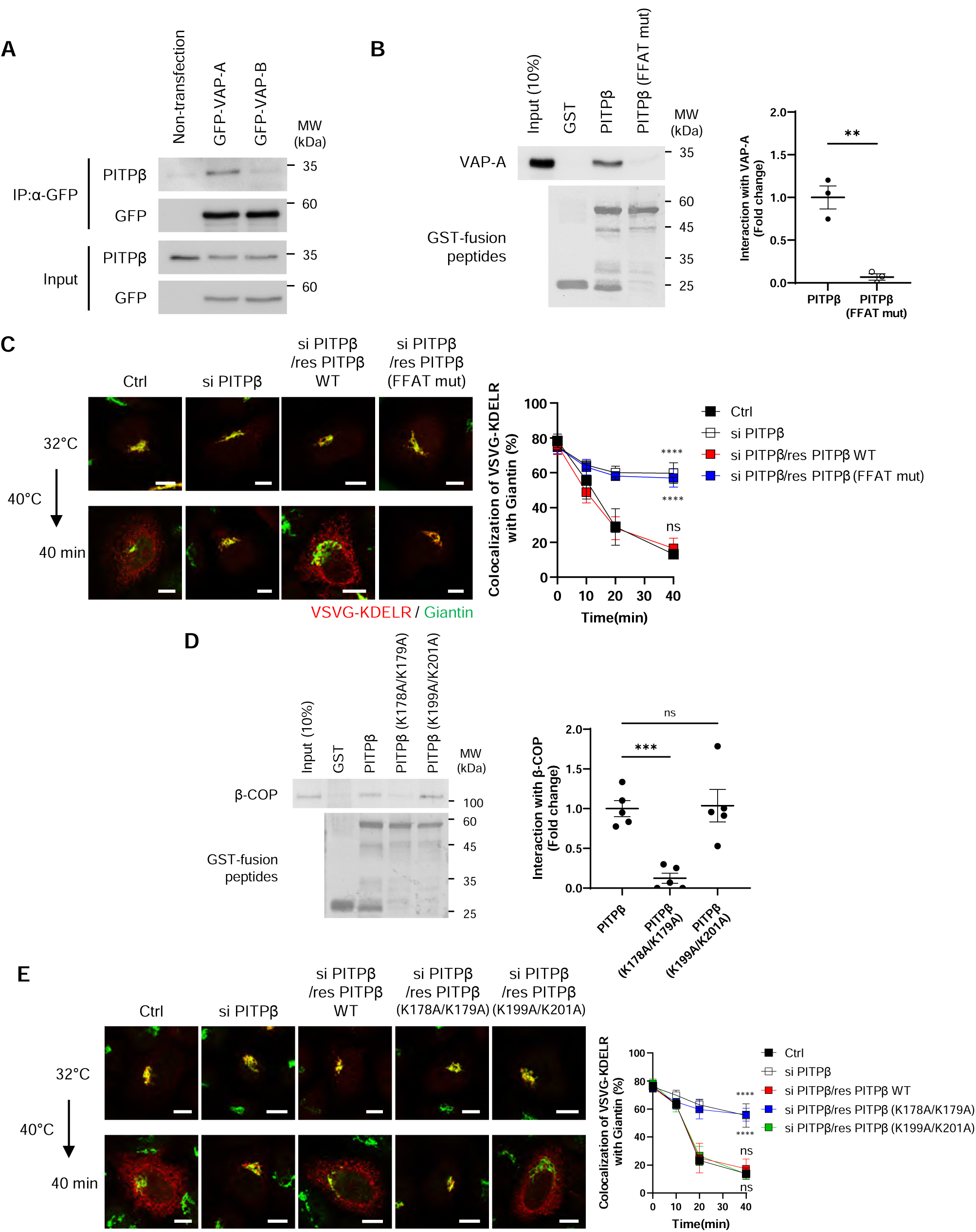
PITPβ interacts with VAP-A and coatomer. Quantitative data are shown as mean ± s.e.m, with the number of independent experiments indicated. Statistics was performed using the two-tailed student t-test: **** p<0.0001, *** p<0.001, ** p<0.01, ns (non-significant) p>0.05. **(A)** Co-precipitation experiment assessing the association of GFP-tagged VAP isoforms with endogenous PITPβ in cells, n=3. **(B)** Pulldown experiment using purified components to assess the effect of mutating FFAT motifs (F107A/F108A) in PITPβ on its direct interaction with VAP-A, n=3. A representative result is shown on left and quantitation is shown on right. **(C)** COPI transport assay assessing the effect of mutating the FFAT motif in PITPβ. Quantitation is shown on right, n=5. Representative confocal images are shown on left, with VSVG-KDELR colored in red and giantin colored in green, bar=10 µm. **(D)** Pulldown experiment using purified components to confirm a specific dilysine motif in full-length PITPβ is critical for its direct interaction with coatomer, n=5. A representative result is shown on left and quantitation is shown on right. **(E)** COPI transport assay assessing the effect of mutating dilysine sequences in PITPβ. Quantitation is shown on right, n=5. Representative confocal images are shown on left, with VSVG-KDELR colored in red and giantin colored in green, bar=10 µm.

### PITPβ anchors to COPI vesicle fission site through multiple interactions

We next sought insight into how PITPβ anchors to the Golgi membrane for this role. Led by our finding above that had revealed the ER to contact the Golgi at sites of COPI vesicle formation, we screened among the COPI-related components by performing pull down assays using purified proteins. We found that PITPβ could interact directly with coatomer (Fig. S3A), but not with ARF1 (Fig. S3B), ARFGAP1 (Fig. S3C), or BARS (Fig. S3D). To confirm the functional relevance of the detected interaction, we next sought to identify specific residues in PITPβ needed for binding to coatomer. Initially, we generated truncations and identified a region in PITPβ that interacts directly with coatomer (Fig. S3E). As this region contains three di-lysine sequences (Fig. S3E), and coatomer is known to recognize di-lysine sequences (Ma and Goldberg, 2013), we next mutated the three candidate sites and identified a single site in the truncated PITPβ that is critical for its direct interaction with coatomer (Fig. S3F). We also confirmed that mutating this site prevents the full-length PITPβ from binding directly to coatomer (Fig. 4D). Subsequently, we expressed this mutant in cells, which involved siRNA against PITPβ followed by rescue using the mutant form, and confirmed that COPI transport was inhibited (Fig. 4E).

### Contact formation likely occurs during COPI vesicle fission

We next pursued another line of investigation that shed further insight into how PITPβ participates in membrane contact. We had found previously that short PA and short DAG are needed sequentially for COPI vesicle fission, which involves PLD2 converting sPC to sPA and then LPP3 converting sPA to sDAG (Park et al., 2019). Thus, as PITPβ delivers the precursor of these critical short lipids, we explored the temporal possibility that PITPβ could be acting in membrane contact more specifically during COPI vesicle fission. We initially performed co-precipitation experiments and found that PITPβ interacts with PLD2 in cells (Fig. 5A). In contrast, PITPβ did not interact with LPP3 (Fig. 5B). We also found that PITPβ mutants (FFAT mutant or K178A/K179A mutant), when transfected and expressed at physiologic level, retained interaction with PLD2 (Fig. S4A and S4B), and thus implicating the interaction with PLD2 to involve a different region of PITPβ than those involved in binding to VAP-A or coatomer.

**Fig. 5.**
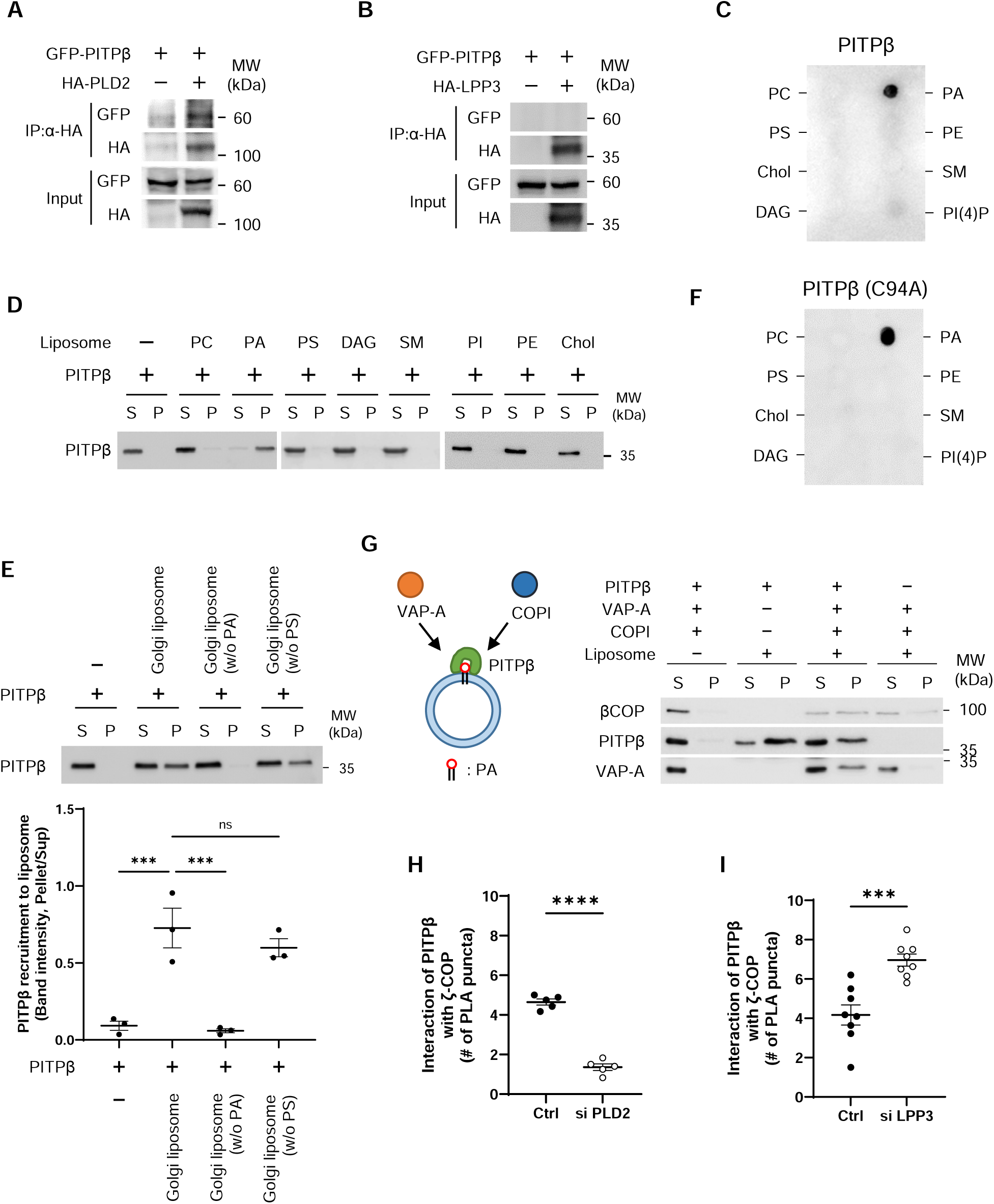
Characterizing additional PITPβ interactions. Quantitative data are shown as mean ± s.e.m, with the number of independent experiments indicated. Statistics was performed using the two-tailed student t-test: **** p<0.0001, *** p<0.001, ns (non-significant) p>0.05. **(A)** Co-precipitation experiment assessing the association of HA-tagged PLD2 with GFP-tagged PITPβ in cells, n=5. **(B)** Co-precipitation experiment assessing the association of HA-tagged LPP3 with GFP-tagged PITPβ in cells, n=5. **(C)** Dot blot analysis revealing that PITPβ binds directly to PC: PC, phosphatidylcholine; PA, phosphatidic acid; PS, phosphatidylserine; PE, phosphatidylethanolamine; Chol, cholesterol; SM, sphingomyelin; DAG, diacylglycerol; PI(4)P, phosphatidylinositol 4-phosphate, n= 4. **(D)** Liposome binding study involving PITPβ incubated with liposome containing different lipids as indicated, n=5. Pellet (P) fraction contains membrane-bound PITPβ, while supernatant (S) fraction contains soluble PITPβ. **(E)** Liposome binding study involving PITPβ incubated with Golgi-like liposomes, either having the full lipid composition or lacking a particular lipid as indicated, n=3. Pellet (P) fraction contains membrane-bound PITPβ, while supernatant (S) fraction contains soluble PITPβ. A representative result is shown above and quantitation is shown below. **(F)** Dot blot analysis revealing that the catalytic mutant of PITPβ binds directly to PC. **(G)** PITPβ can bind simultaneously to PA-containing liposomes, coatomer, and soluble VAP-A, n=3. **(H)** Proximity ligation assay examining the effect of siRNA against PLD2 on the proximity between PITPβ and ζ−COP, n=4. **(I)** Proximity ligation assay examining the effect of siRNA against LPP3 on the proximity between PITPβ and ζ−COP, n=4.

As PLD2 activity generates PA, we next examined whether PITPβ also interacts with PA. As an initial screen, we performed lipid dot-blot analysis, and found that PITPβ could bind directly PA (Fig. 5C). To confirm that this interaction occurs in the context of membrane, we next generated liposomes and found that PITPβ only bound to liposomes generated using PA (Fig. 5D). Moreover, as PLD2 has been found to be concentrated at the neck of COPI buds (Park et al., 2019), we found that increasing the level of PA in liposomes led to increased binding by PITPβ (Fig. S4C). We then generated liposomes having a Golgi-like composition of lipids and confirmed that PA was also needed for PITPβ to bind these liposomes (Fig. 5E). As control, we found that this binding was not affected when phosphatidylserine, which is another negatively charged phospholipid, was withdrawn from the Golgi-like liposomes (Fig. 5E). Thus, PITPβ can anchor to the Golgi by binding not only to protein components (coatomer and PLD2) but also a lipid component (PA) of the COPI bud.

Insight into the portion of PITPβ that binds PA came from a previous study that found a region containing a di-tryptophan (WW) motif to be involved in membrane binding (Shadan et al., 2008). We further noted that this region is distinct from the catalytic domain (Fig. S4D), which suggests how PITPβ can anchor to the Golgi membrane and also catalyze PC transfer. Supporting that these two functions involve distinct regions of PITPβ, we found that the catalytic-dead mutant of PITPβ can still bind PA (Fig. 5F). The structure of PITPβ further suggested that the region responsible for PA binding would be distinct from those involved in binding to VAP-A and to coatomer (Fig. S4D). Thus, PITPβ could bind simultaneously to PA, coatomer, and VAP-A. As confirmation, we generated a truncated form of VAP-A that could not bind membrane and found that PITPβ could bind to this form along with coatomer, while also being recruited to PA-containing liposomes (Fig. 5G).

We also pursued another line of investigation to further confirm that PITPβ binds Golgi membrane through PA. COPI vesicle fission involves the sequential activities of PLD2 and LPP3, with the former generating PA and the latter consuming PA (Park et al., 2019). Thus, we examined whether targeting against PLD2 (to reduce PA level at sites of COPI vesicle fission) would reduce the localization of PITPβ at COPI sites, and whether targeting against LPP3 (to increase PA level at sites of vesicle fission) would enhance the localization of PITPβ at these sites. Upon siRNA against PLD2 (Fig. S4E), we found by the proximity ligation assay that PITPβ showed reduced proximity to ζ-COP (Fig. 5H). Moreover, upon siRNA against LPP3 (Fig. S4F), PITPβ showed increased proximity to ζ-COP (Fig. 5I). These results not only confirmed the importance of PA for PITPβ to bind Golgi membrane, but also further supported that PITPβ forms membrane contact with COPI buds during the fission stage.

### Reconstituting lipid transfer and contact formation by PITPβ

As cell-based studies cannot rule out indirect effects, we next sought to reconstitute lipid transfer using purified components to confirm directly the roles of PITPβ. As done previously to reconstitute lipid transfer through membrane contact (Gong et al., 2021; Mesmin et al., 2013), we generated two populations of liposomes, in this case with one having an ER-like composition of lipids, and another having a Golgi-like composition. For the ER liposomes, we attached VAP-A onto the membrane using a previous established approach that involves tagging a protein with 6x-histidine for its attachment to liposomes that contain a nickel-bound lipid (Lee et al., 2005). For the Golgi liposomes, we incorporated PA into the liposomal membrane to allow binding by PITPβ. To detect PC transfer from the ER liposomes to the Golgi liposomes, we incorporated NBD-tagged PC and rhodamine-tagged PE into the ER liposomes. When both fluorescence-tagged lipids reside in the ER liposomes, fluorescence quenching prevents their detection through their respective fluorophores. However, when PC is transferred from the ER liposomes to the Golgi liposomes, dequenching occurs, resulting in both lipids becoming detectable through their fluorophores (summarized in Fig. 6A).

**Fig. 6.**
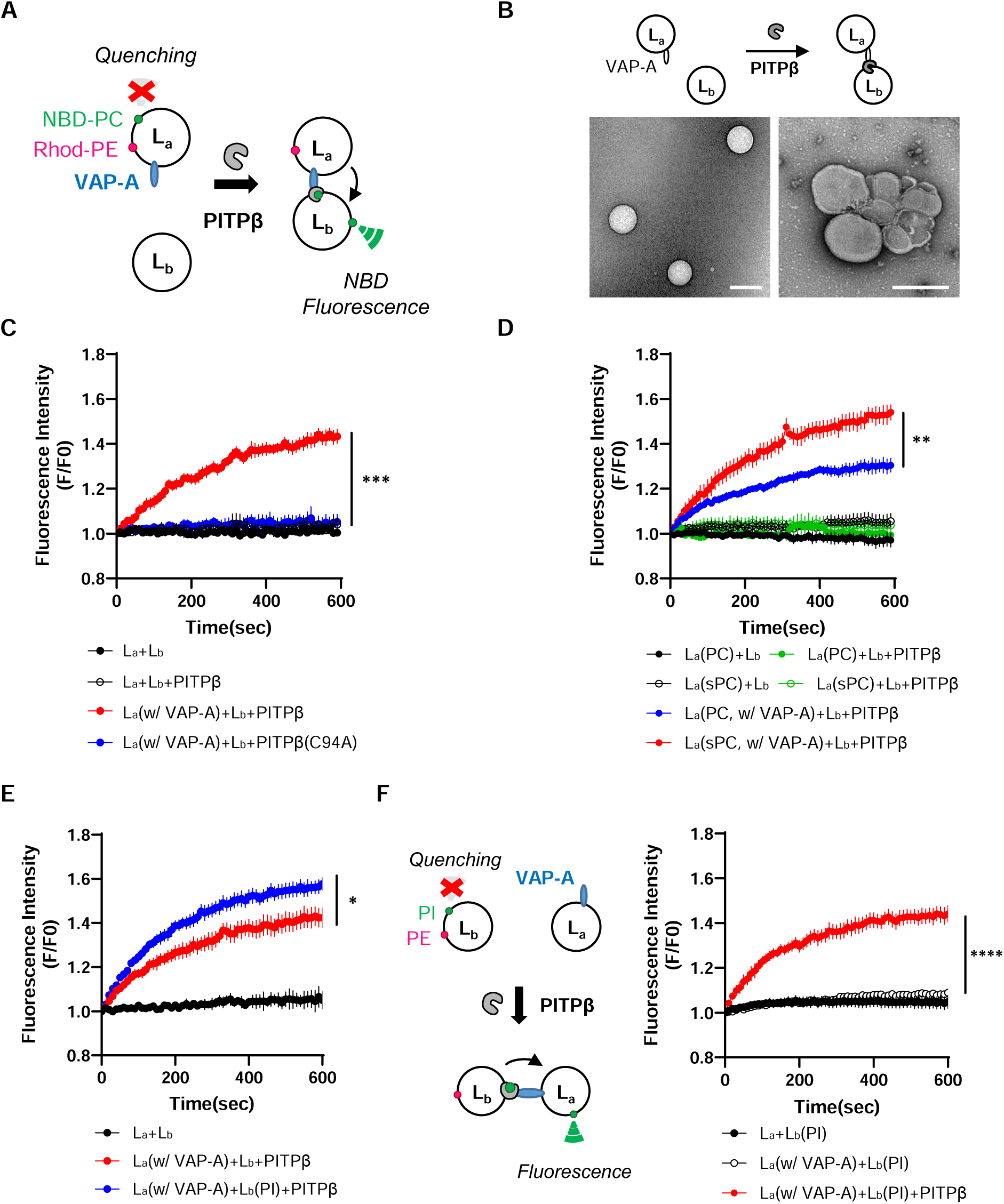
Reconstitution of lipid transfer and contact formation by PITPβ. Quantitative data are shown as mean ± s.e.m, with the number of independent experiments indicated. Statistics was performed using the two-tailed student t-test: **** p<0.0001, *** p<0.001, ** P<0.01, * P<0.05. **(A)** Schematic of the in vitro reconstitution system; L_a_, donor liposome with ER-like composition of lipids. L_b_, acceptor liposome with Golgi-like composition of lipids. 6x his-tagged VAP-A was attached to L_a_ liposomes through nickel-bound lipids. Contact formation were initiated by adding PITPβ and lipid transfer was tracked by measuring the fluorescence signal of NBD-PC over time. **(B)** EM examination confirming the formation of membrane contact among liposomes in the lipid transfer assay, n=4. Representative images of liposomes having contact or not are shown, bar= 200 nm. **(C)** PC transfer requires the catalytic activity of PITPβ, n=5. **(D)** PITPβ preferentially transfers PC with shorter acyl chains, n=5. **(E)** PC transfer by PITPβ is enhanced by incorporating PI into acceptor (L_b_) liposomes, n=5. **(F)** Tracking PI transfer by PITPβ by measuring the fluorescence of Topfluor-PI. Schematics of experimental design is shown on the left. Confirmation of PI transfer from the Golgi liposomes to the ER liposomes is shown on the right, n=6.

We first generated recombinant PITPβ tagged with 6x-histidine to facilitate its purification. We then removed the 6x-histidine tag from recombinant PITPβ after its purification (Fig. S5A) and confirmed that the resulting untagged PITPβ could no longer bind liposomes that contained a nickel-bound lipid (Fig. S5B). Subsequently, when this untagged PITPβ was incubated with ER liposomes and Golgi liposomes, we confirmed contact formation by EM (Fig. 6B), and also that PC transfer was reconstituted (Fig. 6C). As control, transfer did not occur when VAP-A was not attached to the ER liposomes (Fig. 6C). Transfer also did not occur when a catalytic-dead form of PITPβ was used (Fig. 6C). In the latter case, we also ruled out that the defect in PC transfer could be attributed to the mutant protein unable to bind the Golgi liposomes, as it bound to these liposomes similarly as the wild-type form (Fig. S5C). PC transfer also did not occur when PA was not incorporated into the Golgi liposomes (Fig. S5D), or when PITPβ binding to VAP-A was disrupted by using the mutant PITPβ with the FFAT-like sequence mutated (Fig. S5E).

We next found that PITPβ preferentially transfers PC having shorter acyl chains (Fig. 6D). Another insight came from the consideration that cholesterol transfer from the ER to the Golgi had been found previously to involve the counter transfer of PI(4)P (Mesmin et al., 2013). Thus, when also considering that PITPβ transfers not only PC but also phosphatidylinositol (PI) (Allen-Baume et al., 2002), we next incorporated PI into the Golgi liposomes, and found that sPC transfer from the ER liposomes to the Golgi liposomes was enhanced (Fig. 6E). As control, adding PI(4)P to Golgi liposomes did not affect sPC transfer (Fig. S5F). We also confirmed more directly the counter transfer of PI from the Golgi liposomes to the ER liposomes. In this case, we added fluorescence-labeled PI and rhodamine-tagged PE to the Golgi liposomes, and observed dequenching upon the addition of PITPβ (Fig. 6F). Thus, the reconstitution studies not only confirmed the direct roles of PITPβ in catalyzing PC transfer and forming membrane contact, but also revealed new findings that PITPβ preferentially transfers shorter forms of PC, which is enhanced by the counter transfer of PI.

### Confirming PITPβ acts in the fission stage through vesicle reconstitution studies

We also pursued the COPI vesicle reconstitution system to confirm more directly that PITPβ is required for COPI vesicle fission. This approach involves the incubation of Golgi membrane with purified factors, as previously described (Park et al., 2019; Park et al., 2015; Yang et al., 2008; Yang et al., 2018; Yang et al., 2002; Yang et al., 2005; Yang et al., 2011). We initially found that Golgi membrane contained an appreciable level of PITPβ, and high-stringency salt wash could not reduce this level (Fig. 7A). Thus, to collect Golgi membrane depleted of PITPβ, we stably expressed shRNA against PITPβ and then collected Golgi membranes from these cells. Immunoblotting confirmed that Golgi membrane from these cells had reduced PITPβ level (Fig. 7B). We then used this PITPβ-depleted Golgi membrane for the reconstitution system and found that COPI vesicle formation was inhibited (Fig. 7C). Upon EM examination, we observed the accumulation of buds with constricted necks on the Golgi membrane (Fig. 7C). Thus, these results confirmed that PITPβ is critical for the fission stage of COPI vesicle formation.

**Fig. 7.**
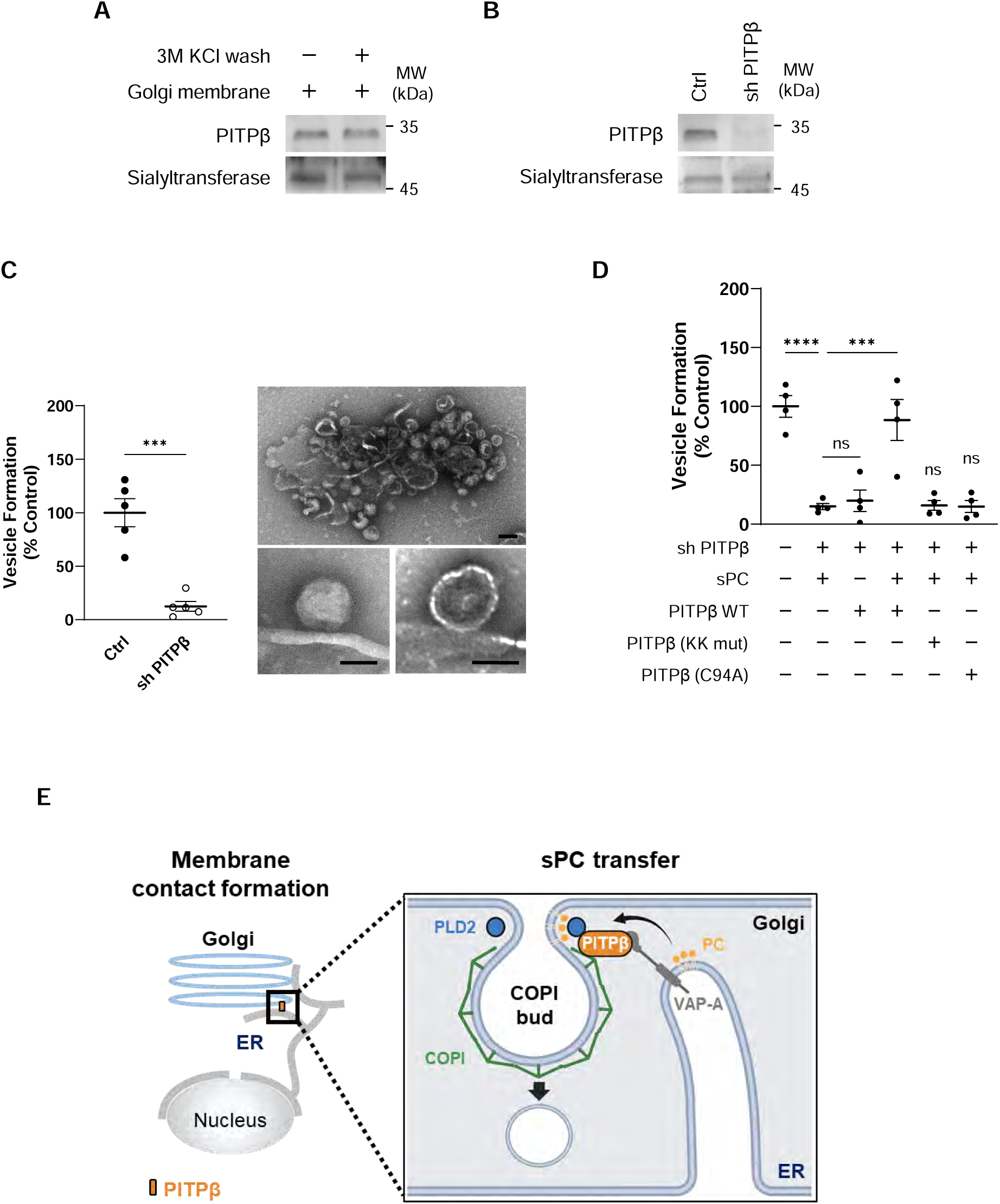
COPI vesicle reconstitution confirming PITPβ acts in vesicle fission. Quantitative data are shown as mean ± s.e.m, with the number of independent experiments indicated. Statistics was performed using the two-tailed student t-test: **** p<0.0001, *** p<0.001, ns p>0.05. **(A)** Western blotting assessing the release of PITPβ from Golgi membrane upon salt wash using 3M KCl, n=4. **(B)** Western blotting assessing the depletion of PITPβ from Golgi membrane by treating cells with shRNA against PITPβ, n=4. **(C)** The COPI vesicle reconstitution system was performed. Left panel shows quantitation of vesicle formation, n=5. Right panels show representative EM images of bud accumulation on the Golgi membrane that had been depleted of PITPβ, bar=50 nm. **(D)** Depletion of PITPβ from Golgi membrane that inhibits COPI vesicle formation is rescued when ER-like liposomes that contain sPC and VAP-A are added along with wild-type PITPβ, but not when mutant forms that cannot interact with coatomer (KK mut) or is defective in catalytic activity (C94A) are added, n=4. **(E)** Schematic summarizing the role of PITPβ in COPI vesicle fission.

We also confirmed that vesicle reconstitution using PITPβ-depleted Golgi membrane requires sPC being delivered to the Golgi membrane. For this delivery, we incubated ER liposomes (having sPC and VAP-A on the membrane surface) and PITPβ with the PITPβ- depleted Golgi membrane (Fig. 7D). As controls, vesicle formation was not reconstituted when incubation lacked either VAP-A coating the ER liposomes or PITPβ as a purified factor (Fig. 7D). Furthermore, when wild-type PITPβ was substituted with mutant forms deficient in binding to coatomer, or catalytic activity, COPI vesicle formation was also not reconstituted (Fig. 7D). Thus, the results confirmed not only that PITPβ acts specifically at the fission stage of COPI vesicle formation, but also its dual role in catalyzing PC transfer and forming membrane contact.

## Discussion

We have identified a specific circumstance that requires the coordination of membrane contact and vesicular transport. Led by our previous finding that sPC is the precursor of short forms of PA and DAG that are needed for COPI vesicle fission (Park et al., 2019), we initially find that sPC is delivered from the ER to the Golgi through membrane contact rather than through COPII transport. We then find that PITPβ plays a key role in this delivery by not only providing the PC transfer activity, but also participating in forming membrane contact (Fig. 7E). As PITPβ is not known to act in membrane contact, our findings achieve a new understanding of how PITPβ acts. Furthermore, whereas PITPβ is known to catalyze both PC and PI transfer, we achieve a new understanding of this role by revealing that both PC and PI transfers are involved in the case of sPC delivery for COPI transport, with PC transfer from the ER to the Golgi enhanced by the counter transfer of PI from the Golgi to the ER.

By combining cell-based studies with reconstitution approaches, we have achieved a high level of spatial and temporal resolution in explaining how sPC is delivered through membrane contact to promote COPI vesicle fission. Spatially, our results reveal an unusual way that a membrane contact can be formed. Rather than two membrane compartments participating in contact formation, a compartmental membrane can also contact a budding carrier for lipid delivery. Temporally, our results suggest that this membrane contact occurs during vesicle fission. Thus, by having sPC being targeted to COPI buds rather than the Golgi membrane more generally, and having this delivery to occur at a specific stage of COPI vesicle formation, our findings reveal how these rare lipids can accumulate to relevant levels locally to support COPI vesicle fission.

At the broader level, we note that membrane contact has been suggested to support vesicular transport, but the precise stage of transport that requires this coordination has been unclear. Our finding addresses this outstanding question by revealing how lipid transfer through membrane contact supports the fission stage of COPI vesicle formation. We further note that COPI transport, being extensively studied, has been a model for elucidating the mechanistic details of vesicle formation. Thus, by advancing the understanding of how COPI vesicle formation occurs, we have also contributed to a basic understanding of how intracellular transport carriers are formed.

## Materials and Methods

### Chemicals, and lipids

The following reagents were obtained: guanine nucleotide triphosphate (GTP) (10106399001) and protease inhibitor cocktail (P2714) from Sigma-Aldrich, H89 dihydrochloride (sc-3537) and Protein G PLUS-agarose (sc-2002) from Santa Cruz Biotechnology, Ni-NTA agarose from Qiagen (30210), and glutathione Sepharose bead from GE healthcare (GE17-0756-01). Lipids were obtained from Avanti Polar Lipid: dioleoyl phosphatidylcholine (DOPC, 850375), dioleoyl phosphatidylethanolamine (DOPE, 850725), dioleoyl phosphatidylserine (DOPS, 840035), dioleoyl phosphatidic acid (DOPA, 840875), Cholesterol (700000), Sphingomyelin (860587), dioleoyl diacylglycerol (DODAG, 800811), dioleoyl phosphatidylinositol (DOPI, 850149), 18:1 phosphatidylinositol-4 phosphate [PI(4)P, 850151], 2C10:0 PC (850325), 2C10:0 PS (840036), 2C10:0 PE (850700), 18:1-12:0 nitrobenzodiazole (NBD)-PC (810133), 14:1-6:0 NBD-PC (810122), 18:1-12:0 NBD-PA (810176), Egg Liss Rhodamine PE (810146), Topfluor-PI (810187), 18:1 DGS-NTA (790404), and 15:0-18:1(d7) PC (791637).

### Proteins, and membranes

Purifications of coatomer, ARF1, ARFGAP1, BARS, and Golgi membrane have been described previously(Fowler et al., 2016). His tagged PITPβ, BARS, and VAP-A (8-212) in pET-15b were expressed in BL21 cells with isopropyl β-D-1-thiogalactopytanoside (IPTG) induction. Cells were lysed in buffer [20 mM Tris-HCl pH 7.4, 200 mM NaCl, 1% Triton X-100, 1 mM dithiothreitol (DTT), and 10% glycerol] containing protease inhibitor cocktail and then subjected to centrifugation. The resulting supernatant was subjected to purification using a Ni-NTA column. PITPβ and proteins fused to glutathione S-transferase (GST) were subcloned into pGEX-4T3 and then expressed in BL21 cells with IPTG induction. Cells were lysed followed by centrifugation, and then the supernatant was subjected to purification using Glutathione Sepharose beads.

### Antibodies

Mouse antibodies against β-COP (M3A5), VSVG (BW8G65), PLD2, and rabbit antibodies against ARF1, ARFGAP1, ζ-COP have been previously described (Park et al., 2019; Park et al., 2015; Yang et al., 2011). The following antibodies were obtained: monoclonal antibody against PITPβ (kind gift from Dr. Shamshad Cockcroft, University College London); anti-HA epitope (3724S), and 6xhis epitope antibodies (9991S) from Cell Signaling; antibodies against GFP (sc-9996), VAP-A (sc-293278), calnexin (sc-46669), and β-actin (sc-47778) from Santa Cruz Biotechnology; antibodies against anti-GM130 (610822) from BD Transduction Laboratories: mouse anti-LPP3 (ab52581) and rabbit anti-giantin (ab80864) antibodies from Abcam; rabbit anti-TGN46 (PA1-1069) and anti-ST6GAL1 (PA5-106647) antibodies from Invitrogen. Secondary antibodies were obtained from Jackson ImmunoResearch Laboratories, and they include horseradish peroxidase (HRP) conjugated Goat Anti-Mouse IgG (H+L) (115-035-003), HRP conjugated Goat Anti-Rabbit IgG (H+L) (111-035-003), HRP conjugated Goat Anti-Mouse IgG Light chain specific(115-035-174), HRP conjugated Goat Anti-Rabbit IgG Light chain specific(211-032-171), Cy3-conjugated Goat anti-Mouse IgG (H+L) (115-165-003), Alexa Fluor 488-conjugated Goat Anti-Rabbit IgG (H+L) (111-545-003), and Alexa Fluor 647-conjugated Goat Anti-Rabbit IgG(H+L) (111-605-003).

### Plasmid, siRNA, and shRNA

Plasmids encoding for PLD2 in pCGN, HA tagged LPP3 in pcDNA3.1, VSVG (ts045) in pcDNA3.1, and VSVG(ts045)-KDELR in pROSE have been described(Gutierrez-Martinez et al., 2013). PITPβ in pEGFP-C1 was a gift from Dr. Shamshad Cockcroft (University College London). PITPβ was subcloned to pET-15b, pcDNA 3.1, and pGEX-4T3 using PCR. Mutant forms of PITPβ were generated using QuickChange II Site-Directed Mutagenesis (Agilent). VAP-A (104447) and VAP-B (104448) in pEGFP-C1 and mEmerald-tagged Sec61β in pEGFP-C1 (90992) were obtained from Addgene. VAP-A(8∼212) was subcloned to pET-15b using PCR. The following siRNA sequences were used against targets: PITPβ (5’-ACGGAUAUUUACAAACUUCCA-3’), VAP-A (5’-CCUGAGAGAUGAAGGUUUA-3’), PLD2 (5′-GGACAACCAAGAAGAAAUA-3′), LPP3 (5′-GGGACUGUCUCGCGUAUCA-3′)(Carvou et al., 2010; Nthiga et al., 2020; Park et al., 2019). Targeting sequence against PITPβ for shRNA expression (5’-CCGGCCATGTTCTGTTCAGGAGTATCTCGAGATACTCCTGAACAGAACATGGTTTTTTG-3’) was synthesized (VectorBuilder).

### Cells

HeLa cells (ATCC), certified as free of mycoplasma contamination, were cultured in Dulbecco’s modified essential medium (Welgene) supplemented with 10% fetal bovine serum (Hyclone), 10 mM HEPES (Gibco) and antibiotics. Transfection of DNA plasmids was performed using GenJet (SignaGen Laboratories). Transfection of siRNA was performed using PepMute (SignaGen Laboratories). Stable expression of shRNA against PITPβ was established by transduction with lentiviral vector followed by puromycin selection.

### In vivo transport assays

Quantitative assays that track COPI and COPII transport in cells have been described previously (Park et al., 2015; Yang et al., 2018). Briefly, HeLa cells were transfected with VSVG(ts045) or VSVG(ts045)-KDELR to track COPII or COPI transport, respectively. For COPII transport, VSVG(ts045) was synchronized at the ER by incubating cells at 40°C for 6 hours and then released by shifting the temperature to 32°C. Cells were fixed at different time points using 4% paraformaldehyde for 10 minutes and then stained using antibodies against giantin (1:500) and VSVG (1:5). For transport at the TGN, VSVG(ts045) was synchronized at the TGN at 20°C for 3 hours and then released by shifting the temperature to 32°C. Cells were fixed at different time points and stained using antibodies against TGN46 (1:200) and VSVG (1:5). For COPI transport, VSVG(ts045)-KDELR was synchronized at the Golgi by incubating cells at 32°C for 6 hours, and then released by shifting the temperature to 40°C. Since 40°C prevents the exit of VSVG(ts045)-KDELR from the ER, this allows one round of retrograde transport from the Golgi to the ER to be assessed. For functional lipid rescue of COPI transport, a lipid of interest (final concentration, 125 μM) was mixed with DOPC (1:1, molar ratio). The lipid mixture was dried using nitrogen gas and then dissolved in 2 ml of pre-warmed culture medium containing bovine serum albumin (BSA) (25 μM). To introduce the lipid to the cells, the lipid-containing medium was incubated with cells for 6 hours before synchronizing VSVG(ts045)-KDELR at the Golgi. Cells were fixed at different time points and then stained using rabbit anti-giantin (1:500) and mouse anti-VSVG (1:5) antibodies. Cells were imaged using Zeiss LSM900 confocal microscopy with Airyscan confocal package containing a Plan-Apochromat 63x objective, Zeiss URGB (488, 561, and 647 nm) laser lines, and Zen 3.8 confocal acquisition software. Images were merged using Image J, and analyzed using MetaMorph 7.7 (MDS Analytical Technologies). Scale bars were generated using Zen 3.8 software.

### Examination of the membrane contacts

Membrane contacts between ER and the Golgi were assessed by two approaches. One approach involves confocal microscopy. Cells were transfected with Emerald-Sec61β using GenJet for 3 days. After fixation using 4% paraformaldehyde, cells were incubated with antibody against giantin to track this cis-Golgi, or against TGN46 to track the TGN, or ζ-COP to track sites of COPI vesicle formation at the Golgi. Imaging used LSM900 Airyscan confocal microscopy followed by quantitation using MetaMorph 7.7 software. Another approach involves the proximity ligation assay (PLA), which used the DUOlink PLA kit (Sigma Aldrich) according to manufacturer’s protocol. Briefly, cells were permeabilized with 0.05% saponin for 20 minutes and then stained using antibody that tracks the ER (calnexin, 1:300), the Golgi (giantin, 1:500), the TGN (TGN46, 1:200), or COPI sites at the Golgi (ζ-COP, 1:300) at 4°C for overnight. Cells were then incubated with oligonucleotide-conjugated secondary antibody at 37°C for 1 hour, followed by incubation with the ligation solution, which contains ligase and circular oligonucleotide, for 30 minutes. The signals of polymerized oligonucleotides were then amplified by incubating the cells with amplification solution containing polymerase at 37°C for 100 minutes. PLA signal was acquired by confocal microscopy and quantified using Image J.

### Immunoprecipitation and GST pulldown

For immunoprecipitation, cells were lysed with lysis buffer (20 mM Tris-HCl pH 7.4, 200 mM NaCl, 1% Triton X-100 and protease inhibitor). After centrifugation at 17,000 x g for 15 minutes, the supernatant fraction was incubated with protein G agarose bead that had been loaded with antibody at 4°C for 2 hours. Beads were rinsed three times using lysis buffer and subjected to SDS-PAGE followed by western blotting. For in vitro pulldown assays, recombinant GST fusion proteins were expressed in bacteria (BL21 cells) using IPTG induction. The cells were lysed using lysis buffer and centrifuged at 4,000 x g for 15 minutes. The supernatant fraction was then incubated with glutathione Sepharose bead (17075601, GE healthcare). GST fusion proteins on beads were incubated with soluble recombinant proteins at 25°C for 2 hours in reaction buffer (25 mM Tris-HCl pH 7.4, 50 mM KCl, 2.5 mM Mg(OAc)_2_, 1 mM DTT, and 0.1% Triton X-100). Beads were washed twice using reaction buffer and then subjected to SDS-PAGE followed by western blotting.

### Lipid dot blot

Purified lipids (200 nmol) were dissolved in chloroform:methanol:50 mM hydrochloric acid (HCl):ponceau S (250:500:200:2, v/v/v/v), spotted on the nitrocellulose membrane (10600002, GE healthcare) and then dried for 1 hour in dark room. After blocking with 3% BSA in TBS (30 mM Tris-HCl pH7.4, 100 mM NaCl) for 2 hours, the membrane was incubated with recombinant PITPβ (1 µg/ml) overnight at 4°C. The membrane was washed three times using TBST buffer (30 mM Tris-HCl pH7.4, 100 mM NaCl, 0.05% Tween-20) and then probed by immunoblotting.

### Liposome binding assay

Liposomes were generated using a mixture of pure lipids. To mimic the Golgi composition, the mixture was: DOPC (50%), DOPE (10%), Cholesterol (17%), Sphingomyelin (8%), DOPS (7%), DOPI (6%), DOPA (1%), and DODAG (1%). To mimic the ER composition, the lipid mixture was: DOPC (70%), DOPE (25%), and DOPS (5%). For simplified liposomes, the mixture was: lipids of interest (30%) and DOPC (70%). Lipid mixture (200 µg) was evaporated using N_2_ gas and then resuspended in 200 µl of reaction buffer (25 mM Tris-HCl pH 7.4, 50 mM KCl, 2.5 mM Mg(OAc)_2_) for hydration at room temperature (RT) overnight and then passed through a 400 nm filter membrane in a mini-extruder (Avanti Polar Lipids). Liposome binding studies were performed by incubating liposomes (20 µg) with 1 µM of soluble recombinant protein of interest in 100 µl of reaction buffer (25 mM Tris-HCl pH 7.4, 50 mM KCl, 2.5 mM Mg(OAc)_2_) at room temperature for 30 minutes, followed by centrifugation at 200,000 x g at 4°C for 30 minutes. The pellet and supernatant fractions were then assessed by SDS-PAGE followed by western blotting to assess the membrane-bound versus the soluble fraction, respectively.

### Electron tomography

A cell slurry (cells resuspended in 20% BSA/PBS) was transferred to B-type planchettes (3 mm) and rapidly frozen with a Leica EM ICE high-pressure freezer (Leica Microsystems). The frozen specimens were freeze-substituted in anhydrous acetone containing 2% osmium tetroxide (OsO_4_) at -80°C for 72 hours in an AFS2 machine (Leica Microsystems). After being slowly warmed to RT over 48 hours, samples were rinsed with 100% acetone three times to remove excess OsO_4_. Cell pellets were separated from planchettes and gradually infiltrated into Embed-812 resin (14120, Electron Microscopy Sciences) in a dilution gradient (10%, 25%, 50%, 75%, 100% in acetone) over 2 days. The samples in 100% resin were polymerized at 60°C overnight. For electron tomography analysis, serial-section ribbons were collected on copper slot grids (150-200 nm) and post-stained. The ribbons were sandwiched between two layers of formvar film and coated with 15 nm gold particles (Ted Pella). Tilt series from ±60 at 1.5° intervals were collected in the brightfield transmission electron microscopy (TEM) mode with a 200-kV Tecnai F20 intermediate voltage electron microscopy (Thermo Fischer). Tilt series around two orthogonal axes were acquired from each section. The IMOD (University of Colorado Boulder) software package was used for tomography reconstruction as described previously(Mai and Kang, 2017; Toyooka and Kang, 2014).

### Reconstitution of COPI vesicle formation

Reconstitution of COPI vesicle formation was performed as previously described with minor modification (Park et al., 2016). Briefly, salt-washed (3 M KCl) Golgi membrane (100 µg) was pelleted by centrifugation at 17,000 x g at 4°C for 30 minutes. The pellet containing Golgi membrane was resuspended using 1 ml of washing buffer (25 mM HEPES-KOH pH 7.4, 3 M KCl, 2.5 mM Mg(OAc)_2_, 1 mg/ml soybean trypsin inhibitor), followed by incubation on ice/water for 5 minutes, and then incubated with myristoylated ARF1 (6 μg/ml), ARFGAP1 (6 μg/ml), BARS (3 μg/ml), coatomer (6 μg/ml), and GTP (2 mM) at 37°C for 30 minutes in 300 μl of reaction buffer (25 mM HEPES-KOH pH 7.4, 50 mM KCl, 2.5 mM Mg(OAc)_2_, 1 mg/ml soybean trypsin inhibitor, 1 mg/ml BSA, and 200 mM sucrose). After the incubation, centrifugation was performed at 17,000 x g at 4°C for 30 minutes to pellet Golgi membrane. Centrifugation was then performed on the supernatant at 200,000 x g using TLA120.2 rotor at 4°C for 30 minutes to pellet vesicles. The level of reconstituted COPI vesicles was determined by western blotting using antibody against β-COP. For PITPβ rescue experiments, recombinant PITPβ (18 μg/ml) was added along with the purified protein factors during the incubation.

### Reconstitution of lipid transfer and contact formation

The reconstitution system was performed similar to that described previously (Gong et al., 2021; Mesmin et al., 2013), with slight modifications. In brief, ER-like (donor) liposomes (L_a_, 64% DOPC, 21% DOPE, 6% DGS-NTA, 5% DOPS, 2% Rhodamine and 2% NBD labeled PC with different chain length; L_a_ (PI), 44% DOPC, 21% DOPE, 20% DOPI, 6% DGS-NTA, 5% DOPS, 2% Rhodamine and 2% NBD-PC) and Golgi-like (acceptor) liposomes (L_b_, 53% DOPC, 18% cholesterol, 11% DOPE, 8% sphingomyelin, 5% DOPS, and 5% DOPA; L_b_ (PI) 33% DOPC, 20% DOPI, 18% cholesterol, 11% DOPE, 8% sphingomyelin, 5% DOPS and 5% DOPA; L_b_ (PI(4)P) 33% DOPC, 20% PI(4)P, 18% cholesterol, 11% DOPE, 8% sphingomyelin, 5% DOPS and 5% DOPA) were prepared. To directly track PI transport from Golgi liposomes to ER liposomes, Golgi-like (donor) liposome (L_b_, 51% DOPC, 18% cholesterol, 9% DOPE, 8% sphingomyelin, 5% DOPS, 5% DOPA, 2% Rhodamine-PE and 2% Topfluor-PI) and ER-like (acceptor) liposome (L_a_, 66% DOPC, 23% DOPE, 6% DGS-NTA and 5% DOPS) was prepared. ER-like liposomes (20 μg) were then incubated with 2 μM of 6x his tagged VAP-A (aa 8-212) at RT for 30 minutes. For lipid transfer and contact formation, donor liposomes (10 µg) were incubated with acceptor liposomes (3 µg) at 37°C in the presence of recombinant PITPβ (1 µM) in which 6x his tag was cleaved. Lipid transfer from donor liposome to acceptor liposome was measured by fluorescence signal of fluorophore-tagged lipid using Synergy H1 microplate reader (Excitation: 463 nm and Emission: 536 nm).

### EM examination of liposomes in the lipid transfer assay

The ER-like liposome (L_a_, 66% DOPC, 21% DOPE, 6% DGS-NTA, 5% DOPS, and 2% Rhodamine-PE), Golgi-like liposome (L_b_, 51% DOPC, 18% cholesterol, 11% DOPE, 8% sphingomyelin, 5% DOPS, 5% DOPA, and 2% NBD-PA) were prepared. ER-like liposomes (10 μg) were then incubated with 2 μM of 6x his tagged VAP-A (aa 8-212) at RT for 30 minutes. For contact formation, VAP-A attached ER-like liposomes were then incubated with Golgi-like liposomes (10 μg) in the presence of recombinant PITPβ (1 μM) for 5 minutes at RT. Liposomes were subsequently imaged using JEOL JEM-1011 electron microscopy.

### Lipid analysis of Golgi membrane by conventional (untargeted) lipidomics

Golgi membrane was extracted using a modified Folch method as previously described (Matyash et al., 2008). Briefly, 200 μl of methanol and 600 μl of Methyl tert-butyl ether (MTBE) containing SPLASH lipidomix internal standard mixture (Avanti Polar Lipids) were added into 350 μg of Golgi sample. After mixing for 1 hour, 200 μl water was added for phase separation followed by centrifugation at 13,000 rpm for 10 minutes. The upper organic phase, MTBE solution containing the lipid extracts, was collected and dried with nitrogen gas. The dried lipid extracts were stored at 4°C and then resuspended in 150 μl of isopropyl alcohol/acetonitrile/water (65:30:5 v/v/v) for UPLC-MS^E^ analysis, which used the same amount of internal standard lipid such as 1-pentadecanoyl-2-oleoyl(d7)-sn-glycero-3-phosphocholine (15:0-18:1(d7) PC) in SPLASH Lipidomix internal standard mixture. The extracted lipids were analyzed using a Waters Synapt G2 HDMS mass spectrometer (Waters), which was operated on the MassLynx 4.1 software and coupled to the AQUITY UPLC system. The chromatographic separation was performed on a CSH C18 AQUITY UPLC column (1.7 μm, 2.1 mm × 100 mm; Waters) with the mobile phases A and B, which consisted of acetonitrile/water (60:40, v/v) and 2-propanol/acetonitrile (90:10, v/v), with 0.1% formic acid and 10 mM ammonium formate. The buffer for mobile phase B was mixed for 3 hours to fully dissolve. The flow rate and injection volume were 0.4 ml/min and 5 μl, respectively. The gradient of mobile phase B was as follows: 0 min, 40%; 2 min, 43%; 2.1 min, 50%; 12 min 54%; 12.1 min, 70%; 18 min, 98%; 18.1 min, 40% and 20 minutes, 40%. The total chromatographic separation runtime was 20 min. We prepared a pooled QC sample at the beginning of the study and aliquots of the same pooled QC sample were used for the entire study. QC samples were measured first and last, and for every six samples. The all ion fragmentation spectra were obtained in positive-ion modes of ESI source. The MS conditions were optimized as follows; i.e., the capillary voltages were 3 kV and 1 kV, the cone voltage was 40 V, the source temperatures were 140°C, and the acquisition mass range of m/z was 50-1200 for positive ionization modes. The high-energy MS spectra were acquired alternatively with the low-energy MS^E^ spectra in UPLC/MSE analysis. The collision energy used for acquiring the high-energy and low-energy MS spectra were 35 and 6 eV, respectively. The high-energy MS spectra provided information on lipid structures. Lipids were annotated using MS-DIAL version 5.1.230429 (Tsugawa et al., 2015). Relative value of each lipid was calculated and visualized as Heatmap using Morpheus software (Broad institute).

### Lipid analysis of sPCs by targeted lipidomics

Golgi membrane (6 mg) was dissolved in 3 ml of methanol:chloroform (2:1, v/v) containing SPLASH Lipidomix internal standard mixture (Avanti Polar Lipids). After adding 1 ml of chloroform and 1.3 ml of 0.1 M of HCl, the mixture was centrifuged at 1,500 x g for 10 minutes. The lower phase containing lipids was collected and evaporated using nitrogen gas, and then reconstituted in methanol chloroform solution (9:1, v/v). PC species were detected by UHPLC coupled electrospray ionization tandem mass spectrometry using an AB Sciex 6500+ Q-trap mass spectrometer system operated in negative ionization/multiple reaction monitoring mode. Lipids were separated by hydrophilic-interaction chromatography using a Luna® 3um NH2, 2 x 100 mm column (Phenomenex Inc) and detected using lipid species specific precursor/product ion pairs. The column was equilibrated using solvent A (93% acetonitrile, 7% dichloromethane and 2 mM ammonium acetate). Samples were eluted using a gradient with solvent A and solvent B (50% acetonitrile, 50% water, 2 mM ammonium acetate pH 8.2) as follows: 0-0.01 min: 0-10% of solvent B, 0.01-9 min: 10-50% of solvent B, 9-10.1 min: 50-100% of solvent B, and 10.1-12.5 min: 100% of solvent B, at a flow rate of 0.6 ml/min. PC species were detected in negative ionization mode as their acetate adducts. Peaks were identified, integrated and reported using AB Sciex OS-Multiquant software. Integrated peak areas for the individual PC species were normalized to the phosphatidylcholine internal standard. Relative value of PC species was calculated and visualized as Heatmap using Morpheus software (Broad institute).

## Acknowledgements

We thank K. Mai for electron microscopy technical advice, and S. Cockcroft for advice and reagents related to PITPβ. This work was funded by grants to S.-Y.P. (National Research Foundation of Korea, 2020R1C1C1008823, RS-2023-00208127, RS-2023-00260454, RS-2024-00344154), to V.W.H. (National Institutes of Health, R37GM058615), to B.-H.K. (Hong Kong Research Grant Council, GRF14113921), and also by BK21 FOUR.

## Author contributions

K.P., S.J. and H.C., performed colocalization studies, lipid transfer studies, immunoblotting, and vesicle reconstitution studies with help from J.J.. K.P., P.G., and B.-H.K. performed EM studies. G. B., J.H.C., and A.J.M. performed lipid analysis. All authors participated in experimental design and data analysis. V.W.H. and S.-Y.P. supervised the project, and wrote the manuscript with input from all other authors.

---

**Fig. S1.**
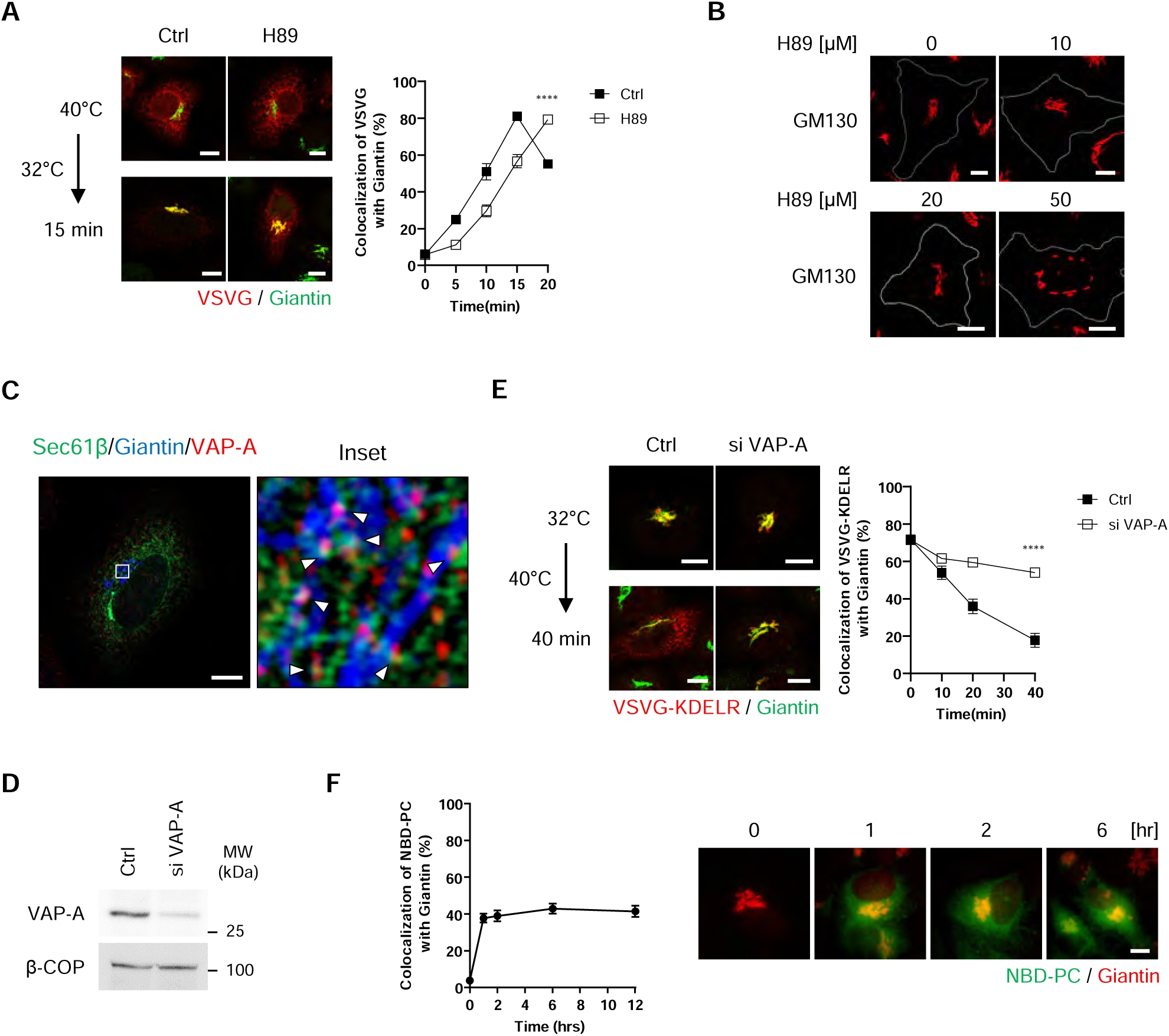
Supporting characterizations of transport and contact formation. Quantitative data are shown as mean ± s.e.m, with the number of independent experiments indicated. Statistics was performed using the two-tailed student *t*-test: **** p<0.0001. **(A)** COPII transport assay assessing the effect of 20 µM of H89 treatment. Quantitation is shown on right, n=3. Representative confocal images are shown on left, VSVG (red) and giantin (green), bar=10 µm. **(B)** Confocal microscopy showing the Golgi remains intact when cells were treated with 20 µM of H89 for 3 hours. Representative images examining the distribution of GM130, a Golgi marker, is shown, bar=10 µm. **(C)** Confocal microscopy assessing the colocalization of VAP-A with an ER marker (Sec61β) and a Golgi marker (giantin), bar=10 µm. **(D)** Efficacy of siRNA against VAP-A as assessed by western blotting of whole cell lysate. β−COP level confirmed similar levels of lysate examined. **(E)** COPI transport assay assessing the effect of siRNA against VAP-A. Quantitation is shown on right, n=4. Representative confocal images are shown on left, VSVG-KDELR (red), giantin (green), bar=10 µm. **(F)** Confirmation of NBD-labeled PC being delivered to the Golgi upon its feeding to cells, as assessed by confocal microscopy. Representative images of NBD-PC (green) colocalizing with giantin (red) are shown on right, bar=10 µm. Quantitation of this colocalization over time is shown on left.

**Fig. S2.**
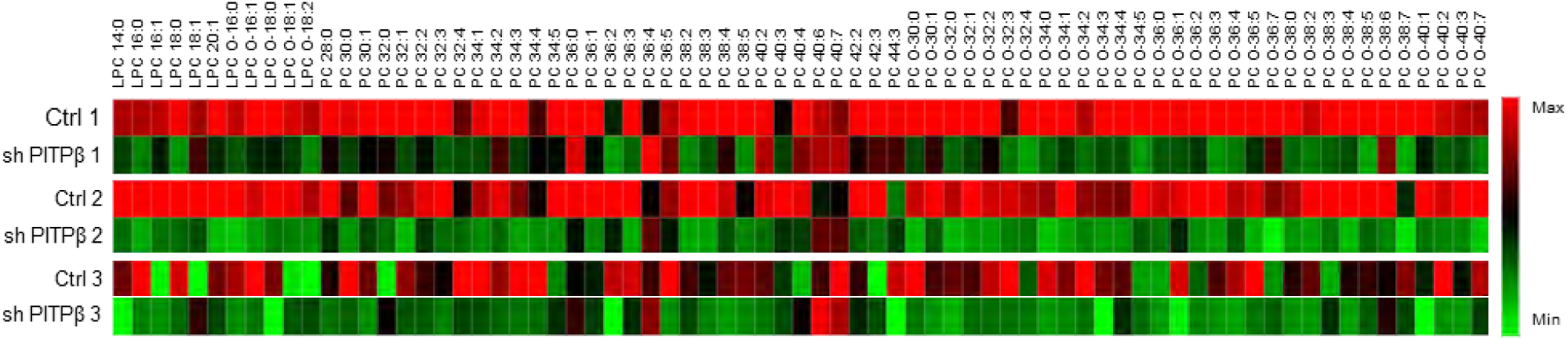
Reduction of PC levels in Golgi membrane upon PITPβ depletion as assessed by untargeted lipidomics, n=3. Results are shown as heat map. PC species are categorized based on their total acyl chain length and unsaturations.

**Fig. S3.**
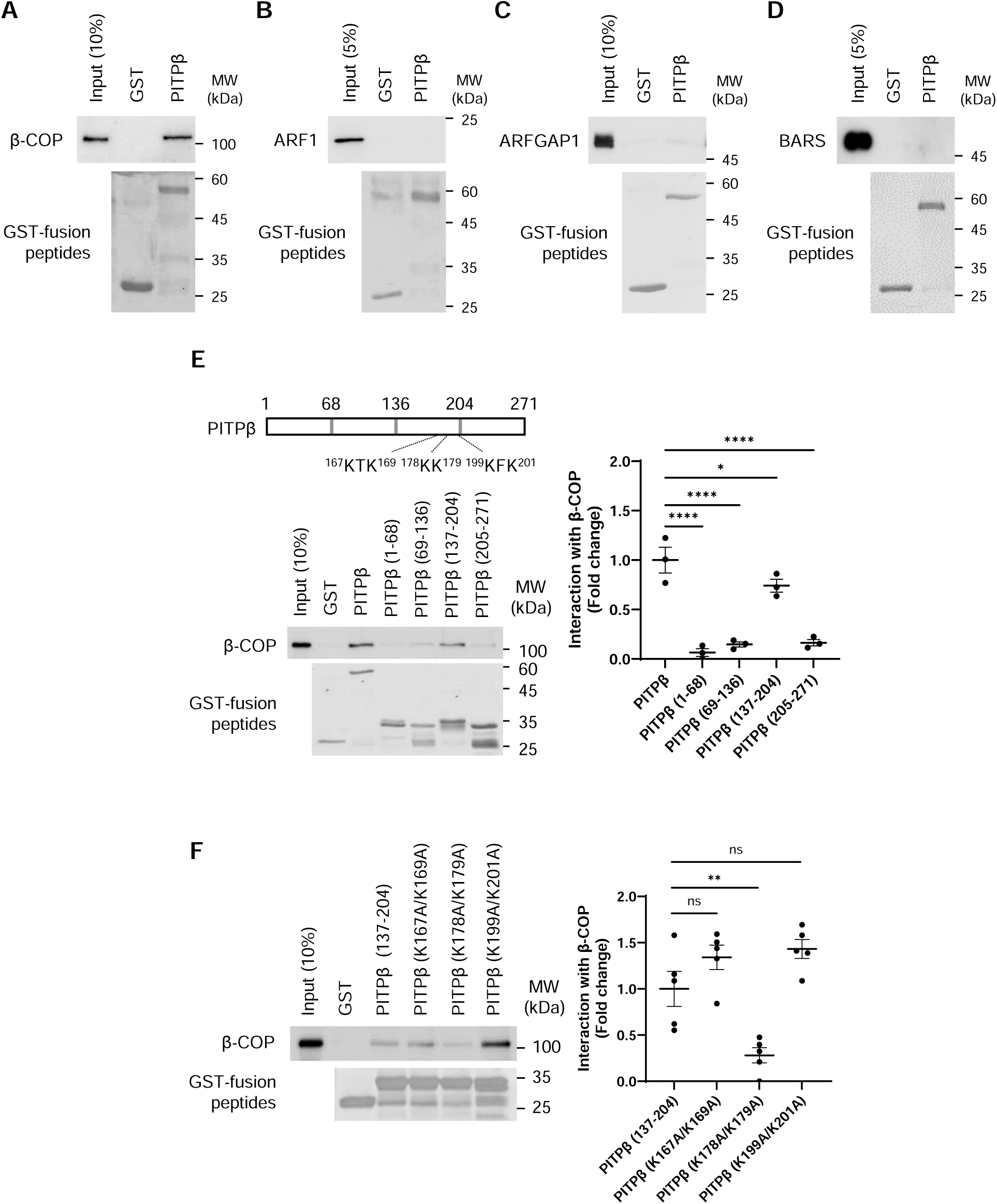
Further characterizing how PITPβ binds to coatomer and VAP-A. Quantitative data are shown as mean ± s.e.m, with the number of independent experiments indicated. Statistics was performed using the two-tailed student t-test: **** p<0.0001, ** p<0.01, * p<0.05, ns p>0.05. **(A-D)** Pulldown experiments using purified components to assess whether PITPβ interacts directly with coatomer (tracked through β-COP), ARF1, ARFGAP1, and BARS, n=5. **(E)** Pulldown experiment using purified components to identify the region in PITPβ critical for its direct interaction with coatomer, n=3. A representative result is shown on left and quantitation is shown on right. **(F)** Pulldown experiment using purified components to identify a specific dilysine motif in truncated PITPβ critical for its direct interaction with coatomer, n=5. A representative result is shown on left and quantitation is shown on right.

**Fig. S4.**
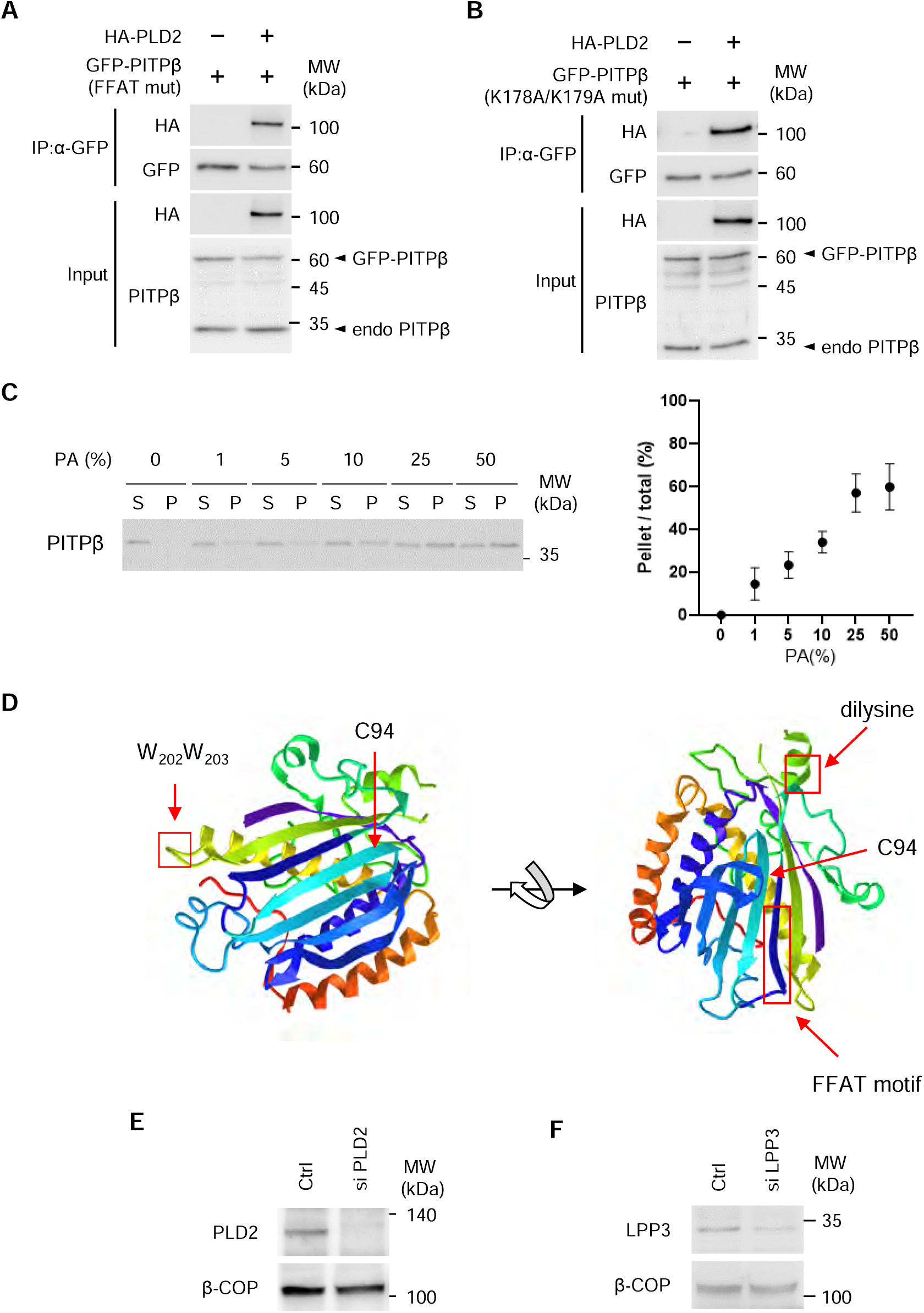
Further characterizing additional interactions of PITPβ. **(A)** Co-precipitation assay showing the association of GFP-tagged PITPβ (FFAA mut) with HA-tagged PLD2, n=4. **(B)** Co-precipitation assay showing the association of GFP-tagged PITPβ (K178A/K179A) with HA-tagged PLD2, n=4. **(C)** PITPβ binding to liposomes that contain increasing level of PA, n=3. A representative result is shown on left, and quantitation is shown on right. **(D)** Highlighting regions in PITPβ responsible for binding to coatomer (dilysine motif), to VAP-A (FFAT motif), and to membrane (WW motif), as well as the catalytic domain (C94). **(E)** Efficacy of siRNA against PLD2 assessed by western blotting of whole cell lysate, with β−COP level confirming similar levels of lysate examined, n=4. **(F)** Efficacy of siRNA against LPP3 assessed by western blotting of whole cell lysate, with β−COP level confirming similar levels of lysate examined, n=4.

**Fig. S5.**
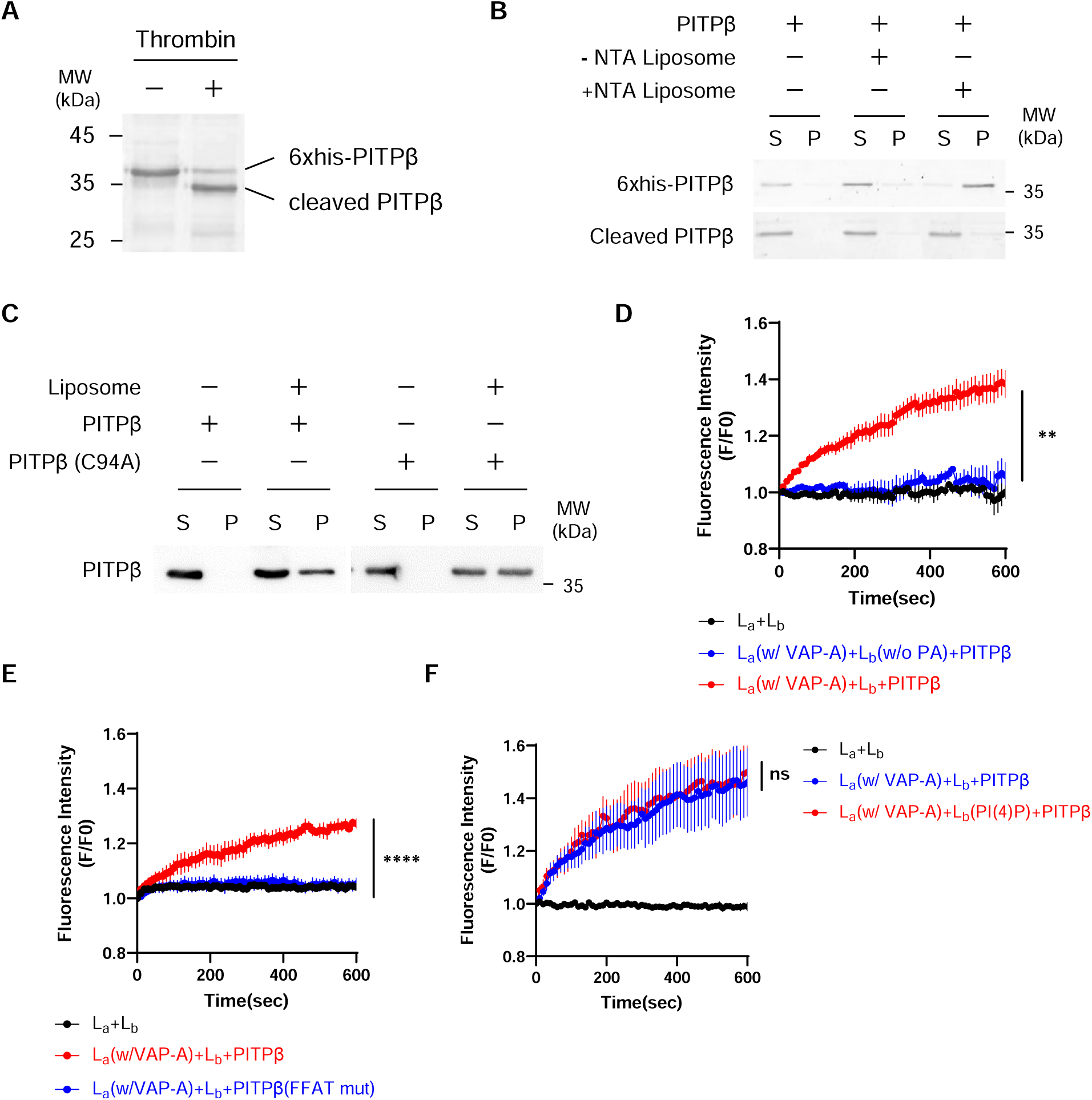
Further characterizing lipid transfer and contact formation by PITPβ. Quantitative data are shown as mean ± s.e.m, with the number of independent experiments indicated. Statistics was performed using the two-tailed student *t*-test: **** p<0.0001, ** p<0.01, ns p>0.05. **(A)** Thrombin-mediated cleavage of the 6x histidine tag from recombinant PITPβ, n=3. **(B)** Liposome binding study examining forms of PITPβ, as indicated, incubated with forms of liposomes, as indicated, n=3. Pellet (P) fraction contains PITPβ on membrane, while supernatant (S) fraction contains soluble PITPβ. **(C)** Liposome binding study involving PITPβ forms, as indicated, incubated with Golgi-like liposomes, n=4. Pellet (P) fraction contains PITPβ on membrane, while supernatant (S) fraction contains soluble PITPβ. **(D)** Reconstitution of lipid transfer and contact formation revealing that PC transfer by PITPβ requires PA incorporated into acceptor (L_b_) liposomes, n=5. **(E)** Reconstitution of lipid transfer and contact formation revealing that PC transfer by PITPβ requires its FFAT-like motif, n=3. **(F)** Reconstitution of lipid transfer and contact formation revealing that PC transfer by PITPβ is not affected by PI(4)P incorporated into acceptor (L_b_) liposomes, n=4.

